# Climate-driven range shifts in fragmented ecosystems

**DOI:** 10.1101/090852

**Authors:** Robin Cristofari, Xiaoming Liu, Francesco Bonadonna, Yves Cherel, Pierre Pistorius, Yvon Le Maho, Virginie Raybaud, Nils Chr Stenseth, Céline Le Bohec, Emiliano Trucchi

## Abstract

Range shift is the primary short-term response of species to rapid climate change but it is hampered by natural or anthropogenic habitat fragmentation. Fragmented habitats expose different critical areas of a species niche to heterogeneous environmental changes resulting in uncoupled effects. Modelling species distribution under complex real-life scenarios and incorporating such uncoupled effects has not been achieved yet. Here we identify the most vulnerable areas and the potential cold refugia of a top-predator with fragmented niche range in the Southern ocean by integrating genomic, ecological and behavioural data with atmospheric and oceanographic models. Our integrative approach constitutes an indispensable example for predicting the effect of global warming on species relying on spatially and ecologically distinct areas to complete their life-cycle (e.g., migratory animals, marine pelagic organisms, central-place foragers) and, in general, on species constrained in fragmented landscapes due to continuously-growing anthropogenic pressure.

## Background

While the impact of anthropogenic climate change on biological communities is beyond question^1^, the nature and extent of species responses are still poorly understood^2^. Species responses to climate change are contingent on intrinsic sensitivity and plasticity^3^ as well as on the poorly-modelled interactions with fragmentation, also human-induced, of species habitats^4,5,6^. The synergy between climate change and habitat fragmentation can pose unforeseen challenges to biodiversity^7,8^, either fragmentation is a natural feature of the ecosystem (e.g. oceanic islands or alpine landscapes) or it is the result of human-mediated land use change^9,10^. Habitat fragmentation is also a peculiar feature of species with complex spatial and temporal distribution of breeding and foraging habitats (e.g. migratory fish, birds and mammals and central-place foragers). In all cases, divergent effects of climate change among distant geographical areas and across trophic levels^11^ impose additional constraints resulting in non-linear responses^12^. Models predicting species response to global warming need to incorporate information on such constraints across the whole species range together with fundamental demographic (e.g. dispersal rate) and trophic parameters^13,14^. Here, we show how a model able to integrate data on habitat distribution, dispersal abilities, population structure, trophic interactions, fitness constraints, and atmospheric and oceanographic scenarios can be used to accurately reconstruct past range shifts in fragmented ecosystems. Such model is applied to forecast demographic and range shift response to current global warming in complex scenarios.

To test our approach, we use a key top-predator of one of the most fast-paced changing ecosystems of our planet, the King penguin (*Aptenodytes patagonicus*), a central-place forager in the sub-Antarctic region and an exemplary case of fragmented distribution of breeding and foraging resources. While a poleward range shift is the predicted response to climate warming for cold-adapted species^15^, the highly fragmented nature of the King penguin’s habitat precludes continuous population displacement. In fact King penguin breeds exclusively on year-round ice-free areas on islands scattered throughout the Southern Ocean and it can only disperse in a stepping-stone manner amongst the few available islands. Its foraging grounds, on the other hand, move together with the myctophid fish stock that strives around the Antarctic Polar Front (APF)^16,17^. The most extensively studied colony, belonging to the most important breeding area for the species (the Crozet archipelago^18^), appears to have benefited from Holocene warming^19^. However, recent tracking studies have revealed a southward extension in their foraging range due to climate change ^17,20^. As a result of the associated increase in energy expenditure related to longer foraging trips, the Crozet population is expected to decline within the coming decades^17,21^. The continuous poleward displacement of the species’ foraging grounds, combined with the discrete distribution of its breeding locations, implies that King penguin populations must undergo abrupt location shifts from island to island to follow their habitat.

### Present and past demographic parameters

In order to predict the limits and opportunities for this species to track its fragmented habitat, we adopted here a cross-disciplinary approach, integrating information from ecology, behaviour and genomics, together with multi-proxy palaeoclimate reconstructions and numerical climate models. The most striking feature of the present-day King penguin population is its worldwide panmixia as recently suggested in both *Aptenodytes* species^22,23^, that we explain by a remarkable migration rate among colonies. Our genome-wide data, including ca. 35,000 independent polymorphic DNA loci genotyped in 163 individuals from 13 different locations covering most of the King penguin’s contemporary range (Extended Data Figure 1; Supplementary Information section S01), strongly contradict the alleged separation between the South Atlantic *patagonicus* and the South Indian and Pacific *halli* subspecies^24,25^, suggesting that the traits used as a basis for subspecies delineation are better explained by phenotypic plasticity than by reproductive isolation. Both classical descriptors of genetic variation and structure analysis unambiguously support a fully-panmictic worldwide population (Supplementary Information section S02). Full admixture among colonies is also clear when repeating these analyses at the island level (Supplementary Information section S02). This result is supported by bio-logging experiments and empirical observations showing short and long distance movements as significant contributors to the ongoing genetic mixing. In addition, new colonies can be established by immigration at a decadal scale^26,27,28^. Contrary to previous hypotheses, recapture of tagged individuals (Supplementary Information section S03) shows that dispersal is also strong at the generation-scale. Thus, dispersal ability is not a limiting factor in the King penguin’s response to environmental change.

The King penguin is a good climatic bio-indicator as confirmed by its strong demographic response to Quaternary climate change. We accurately reconstructed its past demography applying a novel model-flexible approach (the Stairway plot; Supplementary Information section S02.3), based on the composite likelihood of the derived-allele frequency spectrum^29^ calculated on the full high-quality genomic dataset. This analysis was compared to multi-locus Bayesian skyline analyses on a subset of the data (Supplementary Information section S02.4) and pairwise sequentially Markovian coalescent analyses on six additional whole-genome sequences (Supplementary Information section S02.5) and validated through simulations (Supplementary Information section S02.6). The King penguin population experienced two bottlenecks: (a) a recent one during the Last Glacial Maximum (LGM: 19-21 kya), and (b) a more ancient one overlapping with the previous Pleistocene glacial episode (Fig. 1). During the late Pleistocene and early Holocene (ca. 10-17 kya), a period of steep population growth is followed by a long plateau. The large King penguin population fluctuations are not mirrored in the Emperor penguin (Fig. 1) supporting the view that the overall productivity of the Southern Ocean did not change significantly during the Pleistocene and Holocene periods^30,31^.

**Figure 1.**
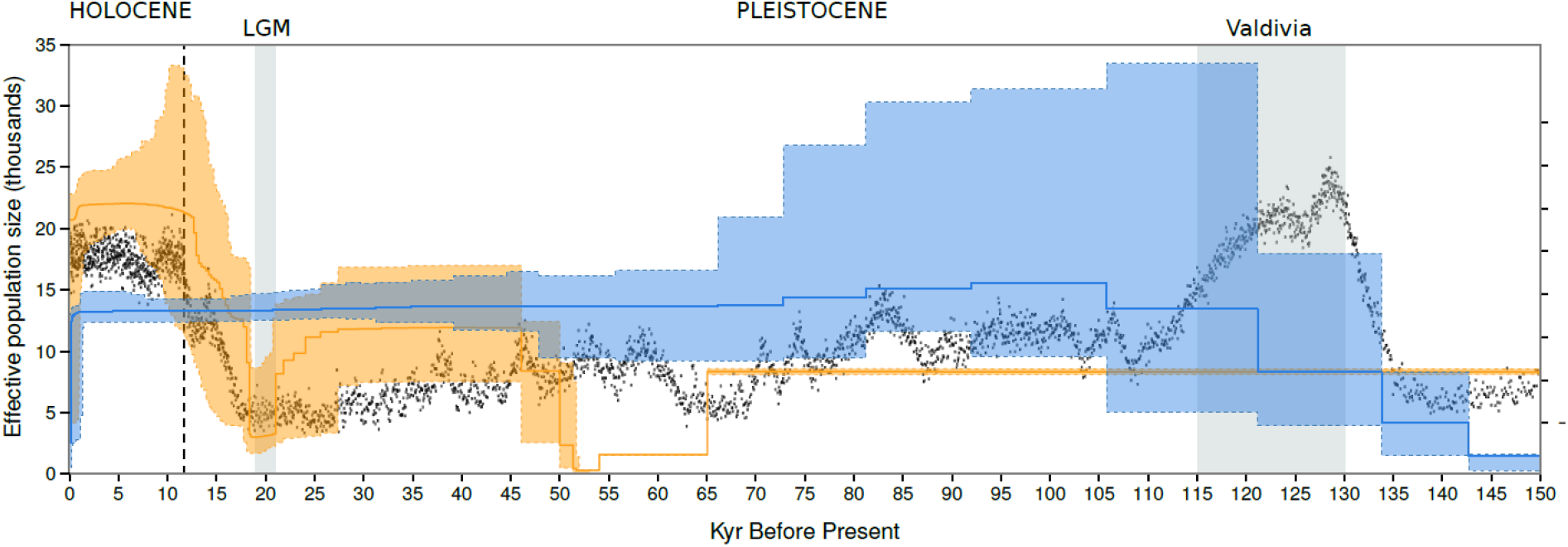
Penguin paleodemography in response to Quaternary climate change. Reconstruction of population size changes (left y-axis) from the last interglacial to present time for the King penguin (orange) and the Emperor penguin (blue). Solid line: median population size; shaded area: 95% confidence interval. Temperature anomaly in the late Quaternary (right y-axis), as inferred from the EPICA Dome C ice core^48^. Highlighted areas: Last Glacial Maximum (LGM, ∼21-19 Kyr BP) and Valdivian interglacial period (∼130-115 Kyr BP). Dashed line: Pleistocene-Holocene transition (∼11.7 Kyr BP). Data for the Emperor penguin are from Cristofari et al^22^.

### Modelling the species range

The King penguin’s response to past climate change is best explained by variations in the extent of suitable habitats (including breeding and foraging grounds; Supplementary Information section S05). We relied on both observed and modelled palaeoclimatic data to identify the extent of the species’ past fundamental niche, which we defined as based on three major traits that directly determine habitat suitability: (a) within foraging distance of the prey stock at the APF^32^, (b) reduced sea ice extent to allow for overwinter chick-rearing^25^, and (c) insular and ice-free land^25^. On the contrary, the location of the APF zone and the extent of land ice and winter sea ice cover exhibited important latitudinal variation over the period^31,33,34^. As a consequence, the location of optimal King penguin breeding areas changed vastly between warm and cold conditions. APF and foraging range predictions, based on historical period (1981-2005) experiments from an ensemble of 15 global coupled ocean-atmosphere general circulation models (from the Coupled Model Intercomparison Project, Phase 5 – CMIP5; Supplementary Information section S05), closely matched both observed APF and empirical foraging distances derived from bio-logging experiments (Supplementary Information section S05.1.1).

Our model is able to capture the full present-day range of the King penguin, and our palaeohabitat reconstructions are also in close agreement with the species’ reconstructed demography (Fig. 1 and 2). Under LGM conditions, the equatorward displacement of the APF and increased land and sea ice cover^31,34^ reduced the King penguin’s range to a fraction of its current extent (Fig. 2A), as suggested by the inferred population bottleneck (Fig. 1A). Assuming a 700-km February foraging distance as the upper limit for successful breeding^20^, the only two possible refugia were found in the Falklands, and in the Campbell plateau region, a much reduced range compared to the eight pre-industrial breeding areas^35^. By mid-Holocene (6 kya), on the other hand, the King penguin already occupied most of its pre-industrial range (Fig. 2B-C). The APF occupied a position close to its present-day state at most locations, while all present-day breeding archipelagos (except for South Georgia) were free from sea ice. Land ice receded early on Kerguelen and South Georgia-although it persisted until early Holocene on Crozet and Prince Edward archipelagos^34^. The King penguin rapidly exploited these newly available locations, as suggested by the steep growth and the following plateau in our demographic reconstructions. Thus, the King penguin’s response to past climate change strongly supports the idea that modifications in the position of the APF and in the distribution of land and sea ice, by modifying the extent of available habitat, have a major impact on the species’ demographic trajectory.

**Figure 2.**
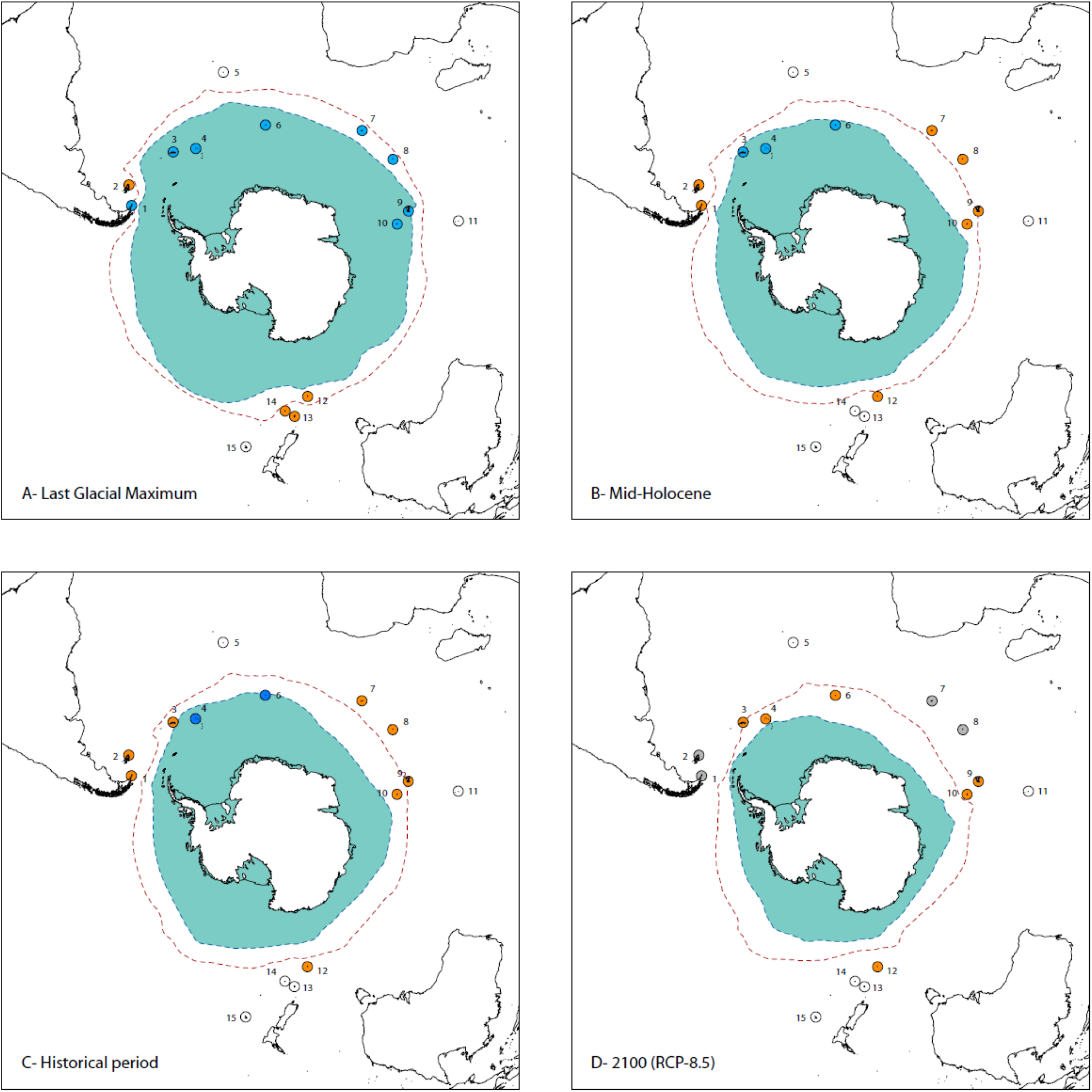
Past and future breeding range of the King penguin. Inferred position of the Antarctic Polar Front, the most important foraging ground for the King penguin, in February (Sea Surface Temperature = 5°C, dashed red line), and extent of sea ice in September (Sea Ice Concentration > 15%, light blue area) at four time periods: **A. Last Glacial Maximum** (21-19 Kyr BP), **B. Mid-Holocene** (6 Kyr BP), **C. Historical period** (1981-2005), **D. Projection for 2100** according to the worst-case greenhouse gas concentration trajectory (Representative Concentration Pathways of +8.5 Watt/m^2^). Occupation status of the islands: **orange:** presence of King penguin breeding colonies, **blue:** sea and/or land ice preventing colony foundation, **grey:** too far from the Antarctic Polar Front for foraging, **white:** never occupied by King penguins. **Islands:** 1: Tierra del Fuego, 2: Falklands, 3: South Georgia, 4: South Sandwich, 5: Gough, 6: Bouvet, 7: Marion and Prince Edward, 8: Crozet, 9: Kerguelen, 10: Heard and McDonald, 11: Amsterdam, 12: Macquarie, 13: Auckland, 14: Campbell, 15: Chatham.

### Forecasting future response to global warming

Projected changes for the 21^st^ century are expected to have a deep impact on the King penguin’s range and population size (Supplementary Information section S05). The uncoupled trends in *(i)* the mobile food resources of the APF and *(ii)* the static breeding locations may have opposite effects depending on the initial state (Fig. 3, Extended Data Figure 2-3): foraging distance increases steadily until the end of the century for the world’s largest colonies, located north of the APF (divergent change); conversely, conditions become more favourable on the colder archipelagos south of the APF, with shorter foraging distances and decreased sea ice (convergent change). This trend is consistent across individual models (Fig 3, Extended Data Figure 2-3) and supported by three different greenhouse gas concentration trajectories (Representative Concentration Pathways +2.6 Watt/m^2^ – RCP-2.6, +4.5 Watt/m^2^ – RCP-4.5, and +8.5 Watt/m^2^ – RCP-8.5) forcing scenarios^36^. With its low genetic diversity and long generation time, the species is not expected to undergo rapid adaptive evolution to the new conditions at the northern end of its range^37,38^: local extinction or dispersal, rather than adaptation, is therefore the predicted outcome.

**Figure 3.**
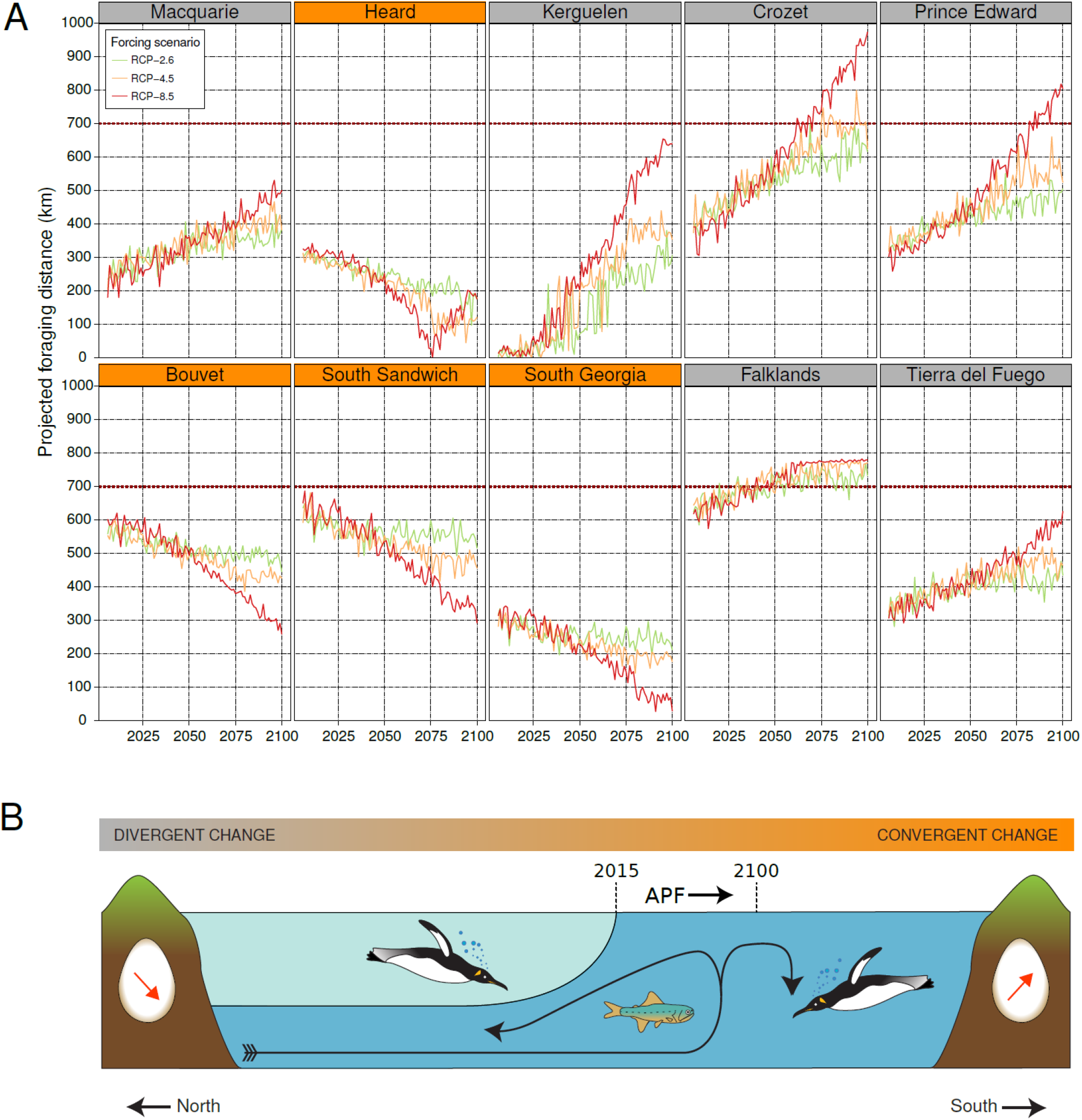
**(A) Projected foraging distance under three greenhouse gas concentration scenarios.** Mean projected summer foraging distance for the King penguin from eight currently occupied archipelagos (see Extended Data Figure 1) and two possible future breeding archipelagos (Bouvet and South Sandwich). Foraging distance are estimated using 15 global coupled ocean-atmosphere general circulation models (from Coupled Model Intercomparison Project, Phase 5-CMIP5), over the 21st century, under three different greenhouse gas concentration trajectories: Representative Concentration Pathways +2.6 Watt/m^2^ (RCP-2.6), +4.5 Watt/m^2^ (RCP-4.5), and +8.5 Watt/m^2^ (RCP-8.5). Horizontal red line: 700 km limit, beyond which no successful breeding of King penguin is expected. Locality name is highlighted according to increasing (gray) or decreasing (orange) distance to the foraging grounds at the Antarctic Polar Front. According to the worst-case scenario (RCP-8.5), King penguin colonies are predicted to (i) disappear from Crozet and Prince Edward, (ii) undergo significant population decline (or disappear) in Kerguelen and newly-colonised Tierra del Fuego, (iii) remain unchanged in Macquarie Island, (iv) grow on South Georgia and Heard, and (v) settle on Bouvet, and possibly the South Sandwich, as the winter sea ice disappears (see Fig. 2). According to low to medium warming scenarios (RCP-2.6 and RCP-4.5, respectively), only Crozet and Prince Edward are too far from the foraging grounds to sustain large breeding colonies by 2100, while Kerguelen retains a favourable situation. **(B) Schematic representation of the different results of climate change in the Southern Ocean.** Dark and light water masses: cold antarctic deep water and warmer subantarctic surface water (major circulation as a black arrow). APF: Antarctic Polar Front. Dashed lines: position of the Antarctic Polar Front in 2015 and 2100 (APF is shifting southward). The red arrow in the egg represents the trend in breeding success. Longer foraging trips from the colony to the APF decrease the breeding success (divergent change, gray) while shorter trips have opposite effect (convergent change, orange).

**Extended Data Figure 1.**
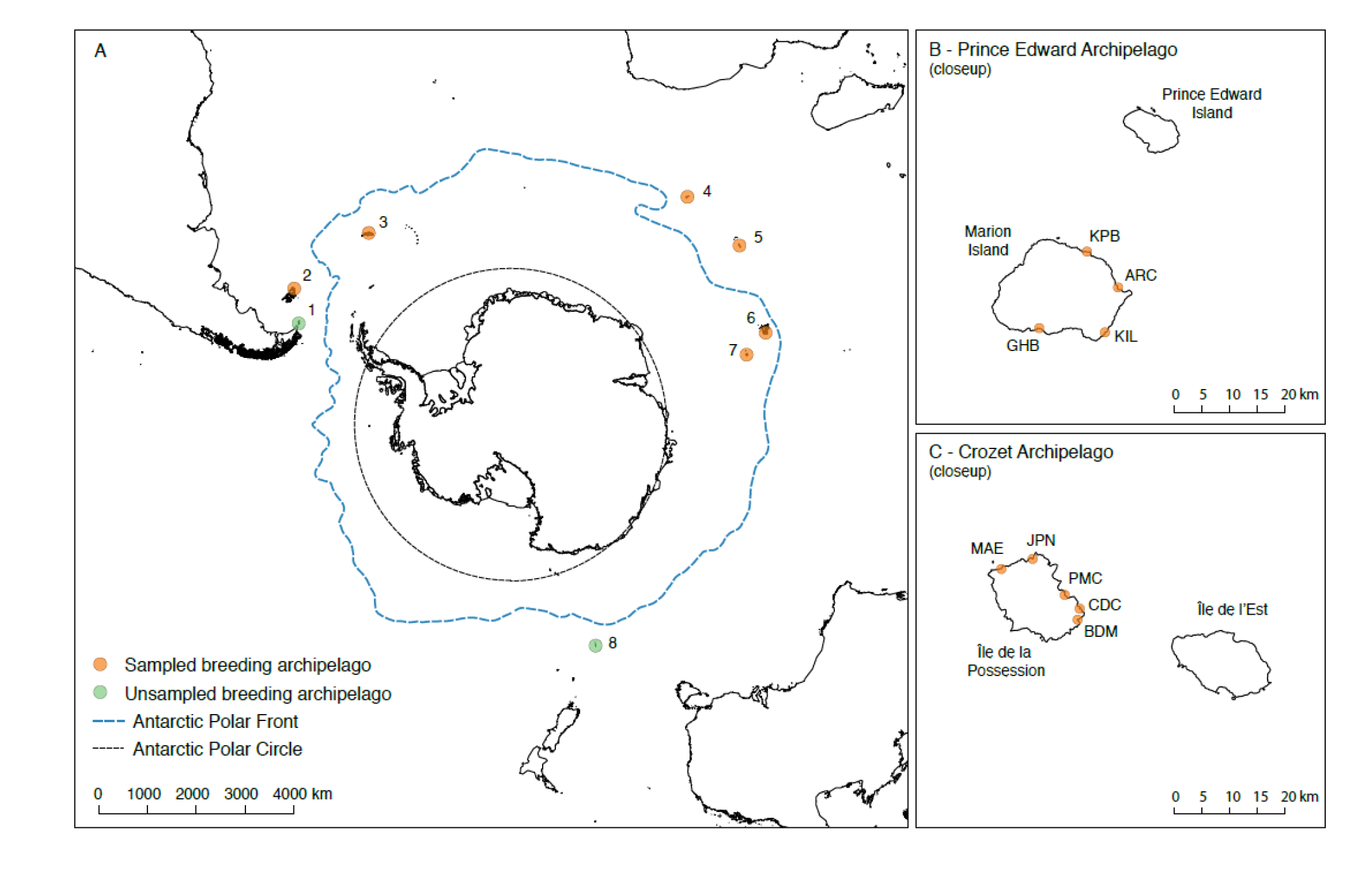
Sampling design. **A)** The King penguin’s range and sampling: (1) Tierra del Fuego, (2) Falklands, (3) South Georgia, (4) Prince Edward archipelago, (5) Crozet archipelago, (6) Kerguelen archipelago, (7) Heard island, (8) Macquarie island. **B)** and **C)** local sampling on Prince Edward and Crozet archipelagos (see Supplementary Information section S01 for details).

**Extended Data Figure 2.**
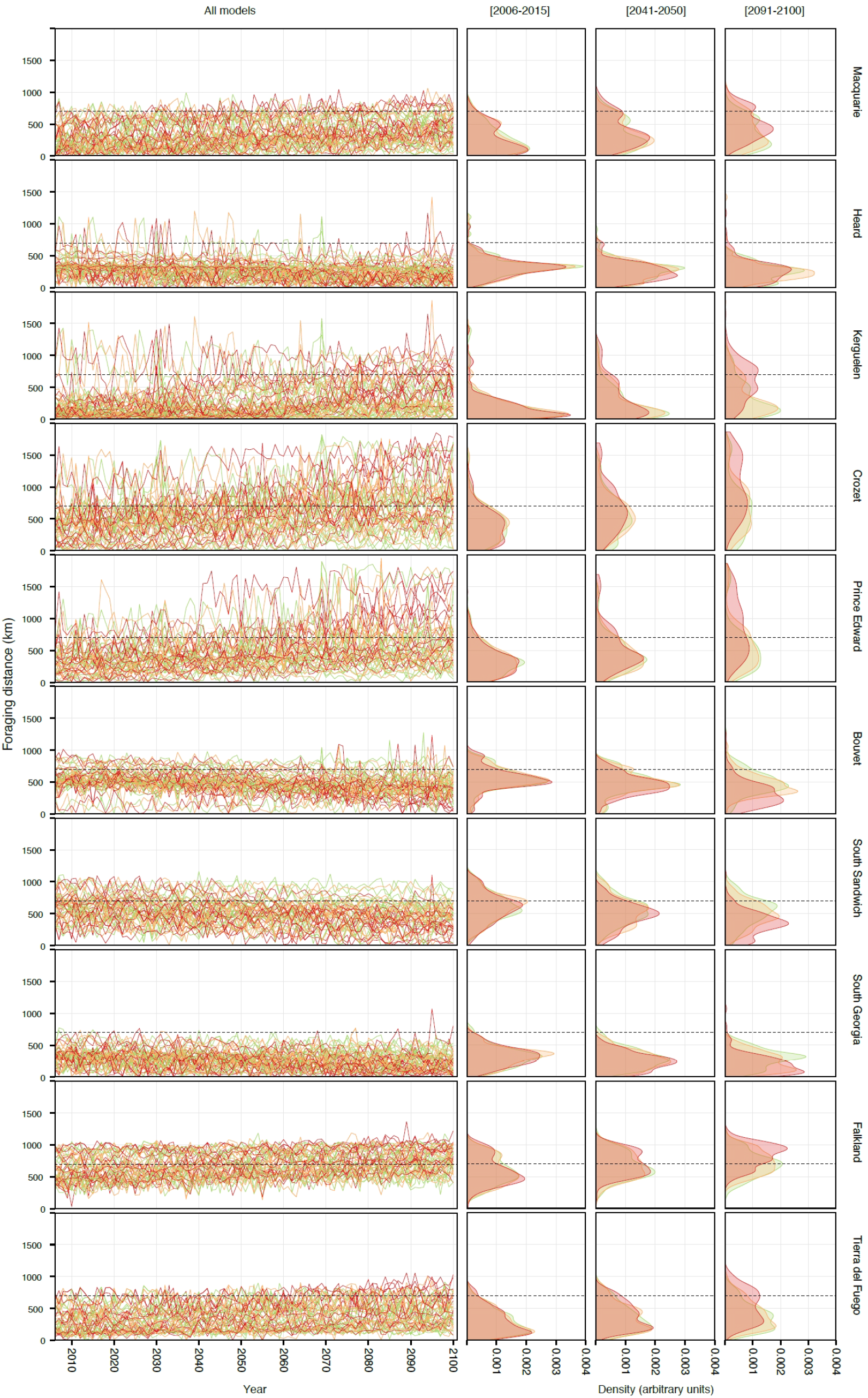
Foraging distance from single models. Projected distance between 10 (eight are currently occupied and two are currently empty but potentially suitable colony locations) subantarctic archipelagos and the Antarctic Polar Front in February estimated from 15 global coupled ocean-atmosphere general circulation models taken separately (from Coupled Model Intercomparison Project, Phase 5-CMIP5), over the 21st century, under three different greenhouse gas concentration trajectories: Representative Concentration Pathways +2.6 Watt/m^2^ (RCP-2.6: green), +4.5 Watt/m^2^ (RCP-4.5: orange), and +8.5 Watt/m^2^ (RCP-8.5: red). Dashed line represents the 700-km limit. Yearly projection (first column of panels); density distribution per RCP scenario, at three different time steps (2nd-4th column of panels).

**Extended Data Figure 3.**
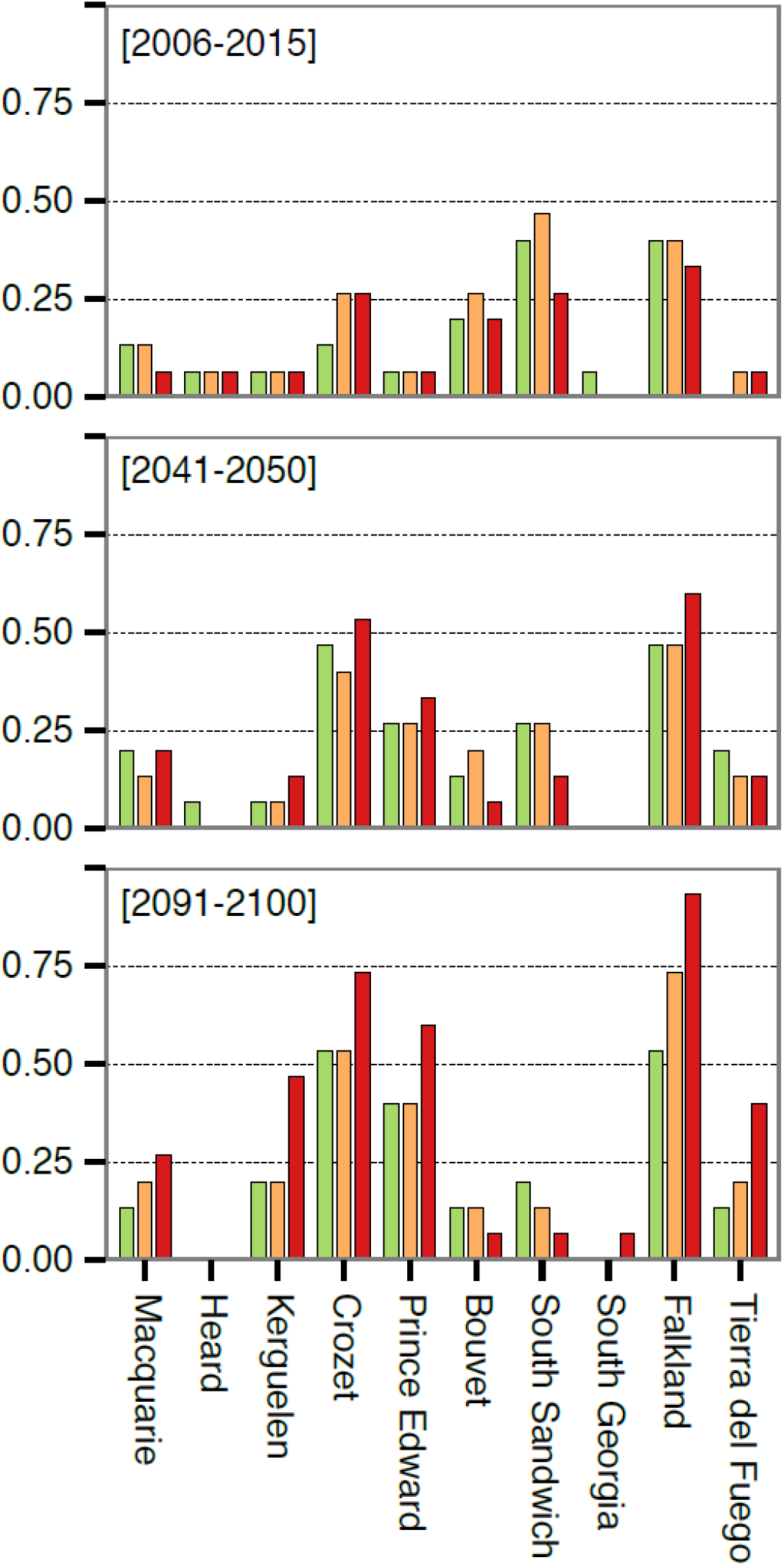
Proportion of models predicting extinction of King penguin colonies. Proportion of the 15 global coupled ocean-atmosphere general circulation models predicting a February foraging distance > 700 km for 20% of the decade, at three different time points. Three different greenhouse gas concentration trajectories are shown: Representative Concentration Pathways +2.6 Watt/m^2^ (RCP-2.6: green), +4.5 Watt/m^2^ (RCP-4.5: orange), and +8.5 Watt/m^2^ (RCP-8.5: red).

Colony loss is likely to bring about a decrease in population size, although high dispersal ability also implies that newly available locations may be colonised rapidly. Under the “*business-as-usual*” RCP-8.5 scenario, 70% of the present-day 1.6 million King penguin breeding pairs^18^ are expected to abruptly relocate or disappear before the end of the century: 49% of the world population are projected to lose their habitat completely (on Crozet and Prince Edward), and 21% will likely see their habitat strongly altered due to regularly near-limit foraging distances (on Kerguelen, Falklands and Tierra del Fuego). These losses may be partly compensated by the predicted colonisation of Bouvet, and by a possible additional growth on Heard and South Georgia due to improved foraging conditions. These last two locations, together with Macquarie Island, are likely to become the major cold refugia for the King penguin in the coming decades. Under the low-emission RCP-2.6 scenario only Crozet and Falkland populations come under direct threat, while other colonies may retain good foraging conditions (Fig. 3, Extended Data Figure 2-3), and undergo minimal demographic impact. Thus, our results stress the importance of immediate action to limit global warming, as efficient attenuation strategies may still have a positive outcome for the Southern Ocean biodiversity. We also insist on the importance of taking proactive conservation measures in areas of the Polar Regions, such as Bouvet, that may act as cold biodiversity refugia for the coming warm-Earth conditions.

Our findings clearly predict a severe disruption in the geographical distribution of this flightless seabird. It is important to note that our projection is likely to be an underestimate, as we only take into account the maximum foraging distance after which no successful breeding may take place. However, increasing foraging distances, even if below the 700 km-limit, have been shown to impact breeding success strongly, and may trigger a colony decrease well before the extinction threshold is reached^17,21^. In addition, our model does not take into account aggravating effects of climate change, such as sea level rise^39^ or decrease in ocean productivity due to ocean acidification^40^ and reduction of the global thermohaline circulation^41^. The abrupt nature of the predicted range shift may also accelerate the restructuring and concentration of biotic interactions (*e.g.* range overlap and competition with other penguin species) generating complex feedback effects not included in our model^38,42^.

## Conclusions

Many species are naturally or artificially constrained in fragmented habitats where the effects of climate change can be enormously exacerbated^43,44,45^. The King penguin’s complex stepping-stone trajectory offers a paradigmatic representation of the impact of global warming on species distributions whenever heterogeneous environmental change leads to uncoupled effects on different critical areas (*e.g.* breeding, foraging, or overwintering grounds). Species distribution modelling is an indispensable tool to foresee the effects of climate change and take preventive measures for biodiversity conservation^46^. Our approach here is the first to take the additional step of integrating uncoupled effects on the whole species range and incorporating information on trophic interactions, population structure, dispersal abilities and past demography, inferred from behavioural and genomic data, with atmospheric and oceanographic models. As a continuously growing number of species are reduced in anthropogenically fragmented landscapes^47^, our integrative method can be extended to all those cases where habitat fragmentation increases the risk of divergent trends in the different portions of a species’ niche, while reducing corridors that may allow continuous niche tracking. By forcing species to undergo tipping point range shifts, habitat fragmentation has the double effect of aggravating the impact of environmental change, but largely masking it, placing populations in a situation of climatic debt well before the critical threshold is reached. Using our approach, we were able to readily identify the most vulnerable areas and to predict the location of potential refugia for a cold-adapted species in a fragmented and rapidly changing environment.

## Acknowledgments

This work was conducted within the framework of the Programme 137 of the Institut Polaire Français Paul-Emile Victor (IPEV; CLB), with additional support from the French National Research Agency (ANR) “PICASO” grant (ANR-2010-BLAN-1728-01; YLM), from Marie Curie Intra European Fellowships (FP7-PEOPLE-IEF-2008, European Commission; project no. 235962 to CLB and FP7-PEOPLE-IEF-2010, European Commission; project no. 252252 to ET), from the Centre Scientifique de Monaco through budget allocated to the Laboratoire International Associé 647 *BioSensib* (CSM/CNRS-University of Strasbourg; CLB, YLM), South African National Antarctic Programme (PP) and the IPEV Programme 109 (YC). Logistic and field costs of research were supported by the IPEV Programme 137 (CLB), the South African Department of Environmental Affairs and National Research Foundation (PP). This work was performed on the Abel Cluster, owned by the University of Oslo and the Norwegian metacenter for High Performance Computing (NOTUR), and operated by the Department for Research Computing at USIT, the University of Oslo. We are very grateful to Morten Skage, Ave Tooming-Klunderud, Marianne Selander-Hansen, and the Norwegian Sequencing Center for their very valuable help in the laboratory, as well as Lex Nederbragt and Michael Matschiner for their assistance with the Abel cluster, and Matteo Fumagalli and Thorfinn Korneliussen for their precious advice regarding ngsTools and ANGSD. We thank Giorgio Bertorelle and Leonida Fusani for their comments on the manuscritp. Genomic data used in analyses are available in the GenBank Sequence Read Archive. We acknowledge the World Climate Research Programme’s Working Group on Coupled Modelling, which is responsible for CMIP, and we thank the climate modeling groups (listed in Table S1 of this paper) for producing and making available their model output. For CMIP the U.S. Department of Energy’s Program for Climate Model Diagnosis and Intercomparison provides coordinating support and led development of software infrastructure in partnership with the Global Organization for Earth System Science Portals.

## Author contribution

CLB and ET conceived and supervised the study. CLB, FB, YC, PP, collected the samples. RC performed DNA extraction, library preparation, and prepared and performed the genomic and demographic analyses and the climate modelling. XL and ET participated in the genomic and demographic analyses. VR and CLB participated in climate modelling. NCS hosted the project. RC, CLB and ET wrote the manuscript. FB, NCS, PP, YC, YLM, VR commented the manuscript.

## S1 Supplementary methods: from sample collection to SNP typing

### S1.1 Sample collection and DNA extraction

A total of 163 blood samples were collected from fledged King penguin juveniles, or from breeding adults, on thirteen colonies covering most of the species’ range (Extended Data Figure 1), between 2010 and 2014. In order to assess fine-scale patterns, we sampled all five colonies from Possession Island, on Crozet archipelago (S46°24’ E51°45’-Baie du Marin “BDM”, N=15, Crique de la Chaloupe “CDC”, N=16, Petite Manchotière “PMC”, N=15, Jardin Japonais “JPN”, N=16, and Mare aux Elephants “MAE”, N=16-fledged juveniles), and all four major colonies from Marion Island (S46°54’ E37°44’-Good Hope Bay “GHB”, N=10, Kildalkey Bay, Archway Bay “ARC”, N=10, and King Penguin Bay “KPB”, N=10-breeding adults). We sampled one colony from Kerguelen archipelago (S49°20’ E69°20’ “KER”, N=16 - fledged juveniles), from Falkland archipelago (S51°45’ W59°00’ - “FLK”, N=10 - all samples were breeding adults), from South Georgia (S54°15’ W36°45’ - “GEO”, N=12 - moulting adults), and from Heard Island (S53°00’ E73°30’ - “HEA”, N=7-breeding adults). Blood was stored in Queen’s lysis buffer at +4°C (Crozet, Marion, Kerguelen), or centrifuged, and red blood cells stored in ethanol at -20°C (Falklands, South Georgia, Heard). DNA was extracted using a spin-column protocol (Qiagen DNEasy© Blood and Tissue kit) with minor modifications.

### S1.2 Genome-wide Single Nucleotide Polymorphism (SNP) typing

SNP discovery and sequencing followed a single-digest RAD-sequencing protocol^1^. Genomic DNA was checked for degradation on a 1.5% agarose gel, and only samples with consistently high molecular weight were retained and quantified by fluorometry (Life technologies” Qubit^®^). Thus, 163 samples were retained and sequenced in 6 distinct libraries. (i) approximately 150 ng of genomic DNA per sample were digested with the restriction enzyme SbfI-HF (NEB); (ii) each sample was then ligated to a unique barcoded P1 adapter prior to pooling in a single library. The library was then sheared by sonication (7 cycles 30″ ON - 30″ OFF); (iii) sonicated libraries were concentrated to 25 μl by DNA capture on magnetic beads (beads solution:DNA = 0.8:1), thus further reducing the carry-over of non-ligated P1 adapters, and the target size range fraction (350-650 bp) was then selected by automated gel electrophoresis (BluePippin^®^); (iv) capture on magnetic beads using the same beads:DNA ratio (0.8:1) was then employed in all following purification steps (after blunt-end repairing, poly-A tailing, P2 adapter ligation and library enrichment by PCR). Magnetic beads were kept together with the library throughout the pre-PCR steps, and DNA was re-bound to the beads for purification using a PEG-8000 binding solution; (v) PCR amplification was performed in 8 × 12.5 μl aliquots pooled after the amplification in order to reduce amplification bias on few loci due to random drift. PCR was performed using NEB Phusion^®^ polymerase with the following cycles: 30" denaturation at 98°C, 18 cycles of amplification (10" at 98°C, 30" at 65°C, and 30" at 72°C), and a final elongation of 5’ at 72°C; (vi) the library was then quantified by a fluorimetry-based method (Life technologies” Qubit^®^), and molarity was checked on an Agilent Bioanalyzer chip (Invitrogen™). A final volume of 20 μl for each library was submitted for paired-end sequencing on an Illumina HiSeq2000 sequencer (V3 chemistry, libraries 1- 3), or HiSeq2500 (V4 chemistry, libraries 4-6), at the Norwegian Sequencing Centre, University of Oslo, spiked with 20% PhiX control library in order to reduce low-diversity bias.

### S1.3 Sequence alignment and genotyping

Data processing was performed using the following workflow: ***(i) Sequence demultiplexing.*** Read quality assessment was made in FastQC (http://www.bioinformatics.babraham.ac.uk/projects/fastqc/). Samples were de-multiplexed according to in-line barcodes using Stacks v1.28^2,3^, low-quality reads were discarded, and sequences trimmed to 95 bp. ***(ii) Read mapping and filtering.*** Demultiplexed fastq files were mapped to the published contigs of the Emperor penguin genome^4^ using Bowtie2 2.2.35, with standard settings, allowing only end-to-end mapping. Resulting SAM files were filtered using Samtools 0.1.196, PicardTools 1.113 (picard.sorceforge.net), and custom R and shell scripts (github.com/rcristofari/RAD-Scripts.git) in order to discard unpaired reads and full read pairs where at least one mate has a mapping quality score below 30. The resulting BAM files were then filtered for PCR and optical duplicates by comparing mapping position and CIGAR string, using Picard MarkDuplicates. This process also allowed to filter out most sequencing errors, since MarkDuplicates only retains the read with the highest average Phred score in each duplicate cluster. ***(iii) SNP calling and genotyping.*** A draft SNP-calling was done in Stacks v1.28 for the general assessment of the dataset, using the “rxStacks” correction algorithm, with a maximum of 5 mismatches allowed between alleles at a single locus (both within and between individuals). For SNP-based analysis, joint SNP and genotype calling was performed using the GATK HaplotypeCaller pipeline^5^, with standard parameters, except for population heterozygosity which was set to 0.01. We retained only SNPs genotyped in at least 75% individuals, or 90% for AMOVA and PCA analyses. ***(iv) Allele-frequency likelihood and allele frequency spectra.*** ANGSD 0.9008^6^ was used to compute per-site probability of being variable, and raw genotype likelihoods, using the Samtools mpileup/bcftools algorithm, and the complete sample allele frequency information as a prior. Per-site allele-frequency likelihood distribution was used to produce a maximum-likelihood estimate of the derived allele frequency spectrum, either unidimensional at the population or species level, or pairwise joint spectrum between pairs of populations.

### S1.4 Ancestral state reconstruction

In order to polarize allele-frequency spectra, we reconstructed the most likely ancestral base for all positions in the RADome. We selected 12 high-quality King penguin samples covering the whole species’ range, and 12 Emperor penguin samples processed according to the same protocol ^7^. We used BEDtools^8^ and GATK’s FastaAlternateReferenceMaker to update the published Emperor penguin genome and establish a reference RADome for both the King penguin, and the Emperor penguin, using only high-quality polymorphisms (phred-scale genotype quality ≥ 80). We aligned this RADome to the Adélie penguin genome (*Pygoscelis adeliæ^9^*) using Bowtie2, and extracted the corresponding regions. For each RAD locus, a maximum-likelihood unrooted tree was built in PhyML^10^, and maximum-likelihood ancestral sequence for crown-Aptenodytes was reconstructed using PAML ^11^ and Lazarus (project-lazarus.googlecode.com/), using PhyML tree topology as a prior. Downstream analysis was restricted to the sites that could be reliably polarized. All sites that were identified as belonging to coding regions ^9^, or to sex chromosomes^12^, were excluded from the analysis.

## S2 Analysis of genetic data

### S2.1 Summary statistics

Summary statistics were calculated in Arlequin^13^ and with custom R scripts either from filtered SNP calls, or from short RAD haplotypes using 11,724 loci. Pairwise fixation index (Fst), calculated using Reich’s estimator^14^, is close to zero (mean pairwise Fst 0.0132 ± 0.00567). Nucleotide diversity π and Tajima’s D were computed for full RAD haplotypes. In order to avoid possible biases due to low coverage, we randomly sampled one haplotype for each individual, and performed calculations on this haploid subset. Tajima’s D is slightly negative, and homogeneous across locations (D_all_: -1.094 ± 0.672, D_HEA_: -0.329 ± 0.925, D_KER_: -0.518 ± 0.899, D_CRO_: -0.546 ± 0.890, D_MAR_: -0.404 ± 0.00307, D_GEO_: -0.448 ± 0.925, D_FLK_: -0.312 ± 0.953), and nucleotide diversity is low (π_ALL_: 0.00209 ± 0.00258, π_HEA_: 0.00201 ± 0.00326, π_KER_: 0.00215 ± 0.00304, π_CRO_: 0.00218 ± 0.00307, π_MAR_: 0.00200 ± 0.00306, π_GEO_: 0.00199 ± 0.00295, π_FLK_: 0.00182 ± 0.00294), in keeping with the prediction of Romiguier *et al.*^15^ for long-lived species.

### S2.2 Descriptive analysis

#### S2.2.1 Principal component analysis

Genotype posterior probabilities calculated in ANGSD (S2.2.1) were used to perform a principal component analysis (PCA) in ngsTools^16^, including 147,711 variable sites with a maximum-likelihood derived allele frequency at least equal to 1/2N (with N being the number of included samples). PCA was repeated in the R package *adegenet*^17^, using a filtered SNP dataset (minimum depth of coverage of 4x, and minimum 80% individuals genotyped at each locus, leaving 4,784 polymorphic loci for analysis). PCA does not resolve strong geographical structure (Fig. S1): although samples tend to gather by archipelago, there is considerable overlap between locations, and no single principal component explains more than ∼0.9% of the total variation.

**Figure S1.**
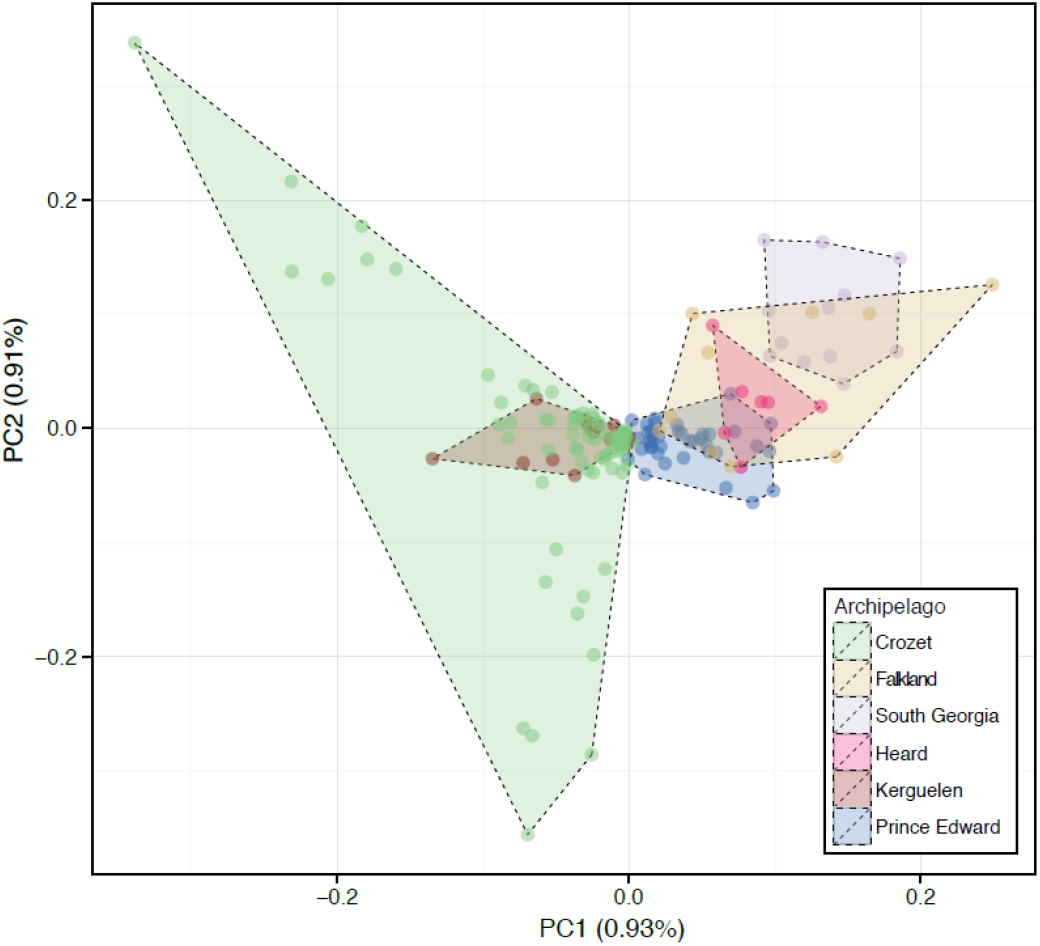
**Principal component analysis** as performed on genotype likelihoods in ngsAdmix^18^, retaining only variable loci. Shaded areas reflect archipelagos.

#### S2.2.2 Clustering analysis

Was performed in ngsAdmix^18^, based on genotype likelihoods calculated in ANGSD with a SAMtools model, and allowing for a maximum of 50% missing data in order to process a site, and keeping only positions inferred as variable with a high likelihood (*p*-value threshold 1e-6). A total of 151,422 sites passed these filters. We performed 100 bootstrap replicates, with K values ranging from 1 to 10. Best-fitting K was chosen using Evanno’s δK method. An independent clustering was performed in FastStructure^19^ using a filtered SNP dataset (minimum depth of coverage of 4x, and minimum 80% individuals genotyped at each locus, leaving 4,784 polymorphic loci for analysis), again with 100 replicates and K ranging from 1 to 10. Both approaches unambiguously supported a K=1 model.

#### S2.2.3 Analysis of molecular variance

Analysis of molecular variance was performed in Arlequin 3.5.2.1^13^, using a filtered SNP set that included only sites genotyped in 90% individuals (62,625 polymorphic sites). Amova was performed on a per-locus basis, with 10,000 permutations. We tested four different grouping schemes: *(i) colonies grouped by archipelago*: ((HEA), (KER), (BDM, CDC, PMC, JPN, MAE), (GHB, KIL, ARC, KPB), (GEO), (FLK)) *(ii) A. p. patagonicus vs A. p. halli*: ((HEA, KER, BDM, CDC, PMC, JPN, MAE, GHB, KIL, ARC, KPB), (GEO, FLK)) *(iii) Crozet-only*: ((BDM), (CDC), (PMC), (JPN), (MAE)) *(iv) Marion-only*: ((GHB), (KIL), (ARC), (KPB)). Under all four groupings, the overwhelming majority of variance is explained at the individual level: *(i)* 92.9% within individuals, 6.20% amongst individuals, 0.989% amongst populations, -0.124% amongst groups. *(ii)* 92.9% within individuals, 6.20% amongst individuals, 0.904% amongst populations, -0.0370% amongst groups. *(iii)* 94.1% within individuals, 4.57% amongst individuals, 1.30% amongst populations. *(iv)* 85.1% within individuals, 14.8% amongst individuals, 0.0309% amongst populations.

#### S2.2.4 Pairwise Hamming distance network

We calculated pairwise Hamming distance between individuals based on genotype calls using PLINK v1.9^20^ using 62,625 polymorphic sites genotyped in 90% individuals, and calculated the corresponding neighbour-net in SplitsTree^21^ (Fig. S2-A). In keeping with the results of AMOVA and PCA, the terminal branches explain most of the variance, and samples do not cluster geographically.

#### S2.2.5 Mitochondrial DNA comparison between Crozet and Macquarie Island colony

Comparison of mitochondrial hypervariable control region (HVR) haplotypes from Crozet (Trucchi et al. ^22^, from 139 individuals, Genbank accession number KF530582-KF530621) with published sequences from Macquarie Island (Heupink et al.^23^, 35 individuals, Genbank accession number JQ256379-JQ256413) confirms the idea of a single, worldwide and fully panmictic population. Pairwise Fst is low (Fst=0.032), and a haplotype network does not support any population separation between the two islands (Fig. S2-B).

**Figure S2.**
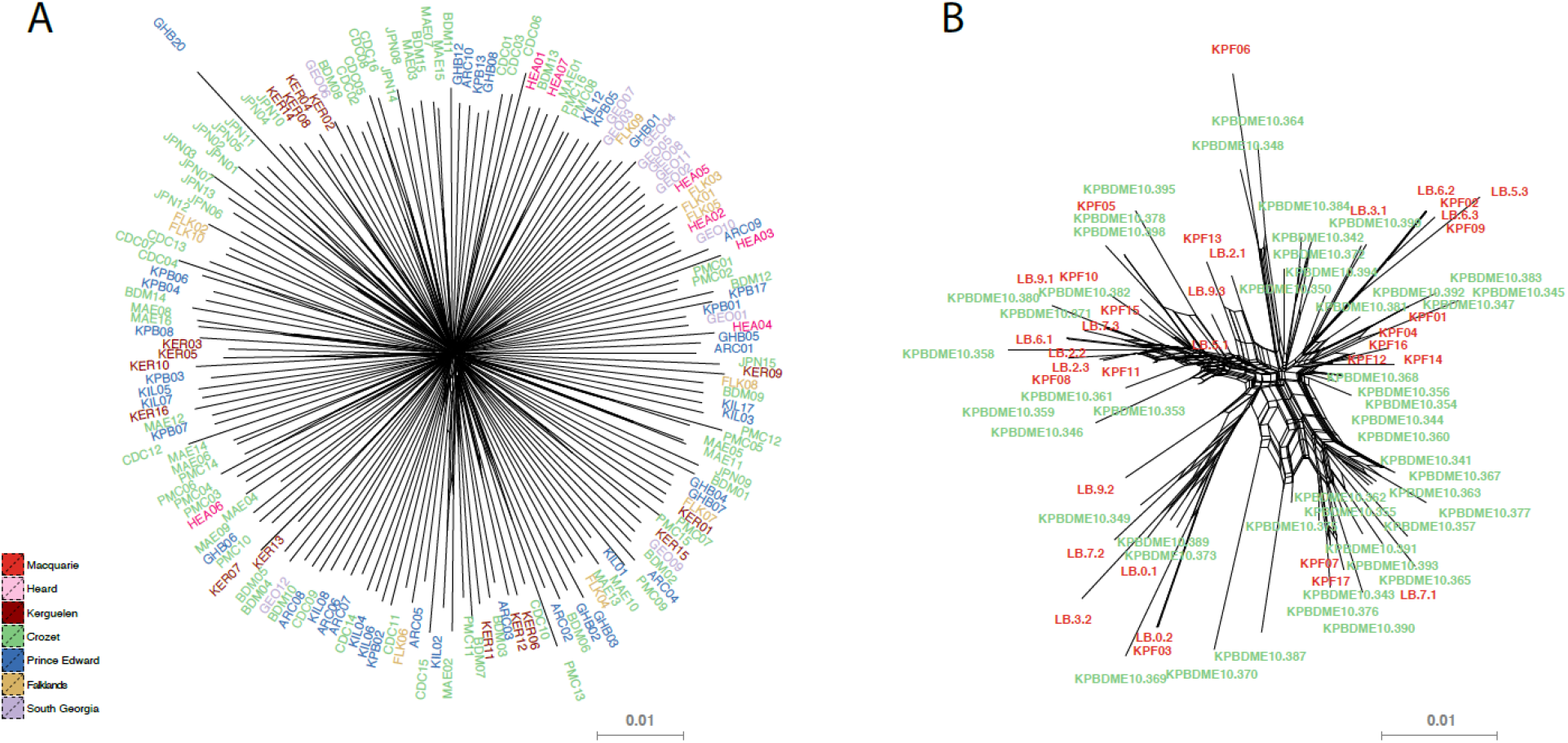
**Neighbour-net** calculated **A)** from pairwise Hamming distances, based on genome-wide SNP data, for 6 breeding archipelagos, and **B)** from the mitochondrial control region of 40 individuals from Crozet, and 39 individuals from Macquarie island.

### S2.3 Demographic reconstructions I: the Stairway plot method

The Stairway Plot method is a novel method for demographic inference developed by Liu & Fu^24^. This model-flexible method relies on the maximisation of the composite likelihood of the observed derived-allele frequency spectrum, without prior hypothesis on demographic history, as opposed to previous spectrum-based demographic inference methods (e.g. Gutenkunst^25^). Maximum-likelihood estimation of the allele frequency spectrum was performed in ANGSD-0.901 under a SAMtools model, for 140 high-quality King penguin samples, and 90 high-quality Emperor-penguin samples using 2,300,996 sites. Each spectrum was run along with 500 bootstrap replicates. Singletons were found to be the least robustly estimated frequency class, due in particular to the confounding effect of sequencing errors, and were consequently masked from the reconstructions - although comparison of reconstructions *(i)* including all frequency categories, *(ii)* excluding singletons, or *(iii)* singletons and doubletons show that only the reconstruction of the most recent demographic events are affected by the low-frequency variants (Fig. S3A-C). Similarly, using only a randomly picked subset of half of the individuals did not affect the reconstructions (Fig. S3D) ***Generation time:*** In a long-lived species, generation time is not a fixed parameter, but rather a function of the demographic trend. An estimator has been defined by Saether et al.^26^ as α + (S / (λ–S)), where α is the age at first breeding for females, S is the yearly adult survival rate, and λ is the yearly growth rate of the population, defined as λ=1 for a stable population. Using long-term monitoring data, we extracted both yearly growth rate, and adult survival, from a pool of 400 adults of known age (Le Bohec *pers. com.*), for the 1999-2010 period. S and λ were found to be strongly correlated over that period (intercept: -0.2454, slope: 1.0936, R^2^=0.6): therefore, we extended the empirical relationship between both parameters to our reconstruction. For each generation, the generation time in years was therefore defined as T*t*+1 = α + (St / (λ*t*–S*t*)), where λt and St are the growth rate and adult survival rate for the previous generation, defined as λ*t* = ( N*t*+1 / N*t*).e(1/T*t*), where N*t*+1 and N*t* are the population sizes at generations *t*+1 and *t*, and T *t* is the generation time in years at generation *t*, and St is a linear function of λ*t*, using empirically derived parameters. This correction was applied recursively from the oldest generation in the reconstruction assuming λ = 1, and towards the present. In order to calibrate other analyses, the mean generation time over the whole reconstruction T = 10.6 years was retained.

**Figure S3.**
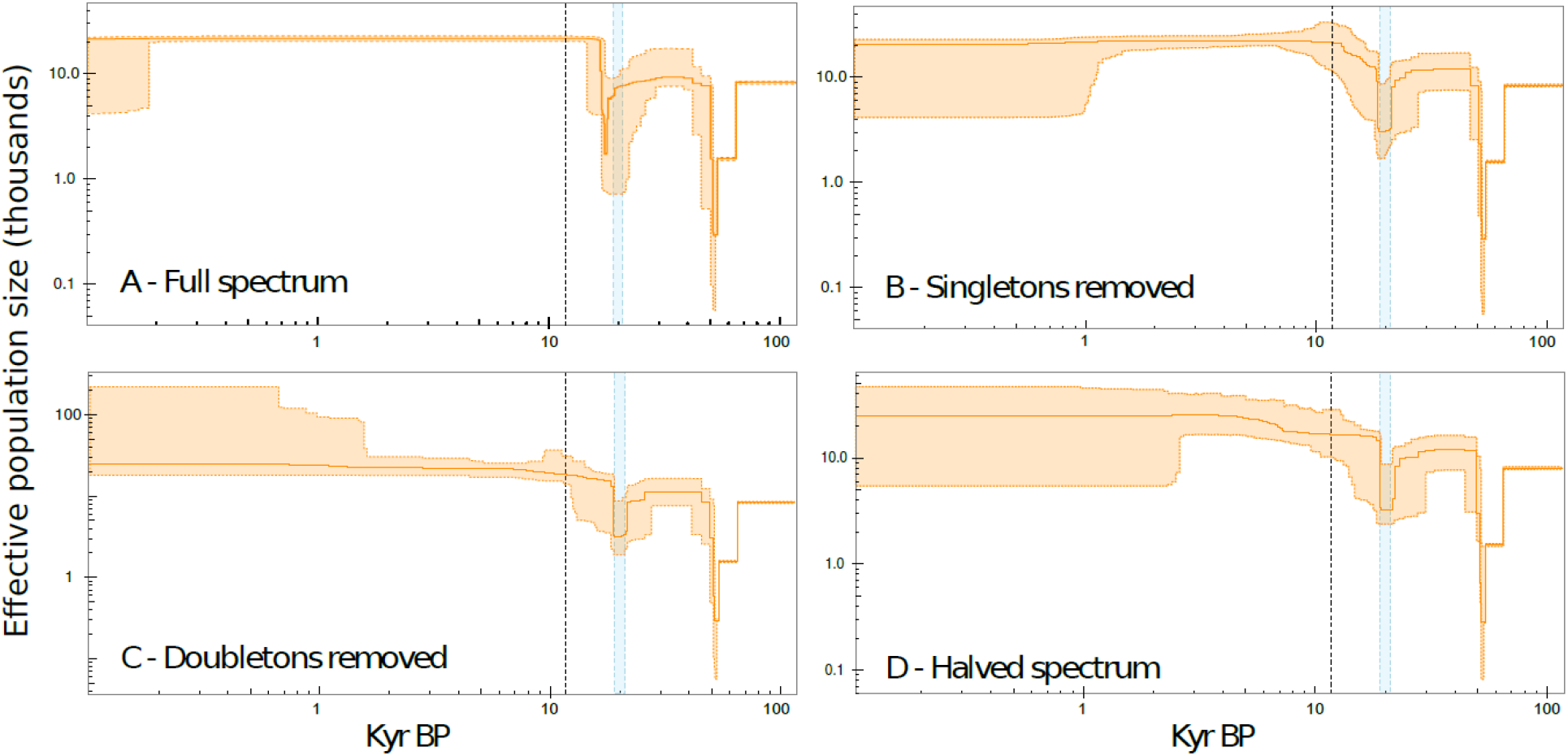
Robustness of the Stairway plot method. Stairway plot reconstructions with the same set of King penguin individuals, but based on **A)** the full spectrum as inferred from 140 samples, **B)** masking the singleton loci, C) masking both singletons and doubletons, and **D)** masking singletons, and using only a random subset of 70 individuals. Dashed black line: Pleistocene-Holocene boundary. Shaded blue band: last glacial maximum. Plots were cut between ca. 100,000 and 100 years before present.

### S2.4 Demographic reconstructions II: the Extended Bayesian Skyline Plot (EBSP) method

Accurate reconstruction of past and present population size changes requires a robust estimate of the substitution rate. We performed a joint analysis of mitochondrial HVR and RAD data, in a multilocus EBSP framework, using the robustly established substitution rate for the Adélie penguin HVR (in substitutions per site per Myr: median = 0.55, 95% CI = 0.29–0.88^27^) as a calibration. Since the generation time differs widely between the Adélie penguin (6.46 years^27^) and the King penguin (10.6 years, see **S2.3**), and since we are considering the rate of substitution as determining the frequency of coalescence events, as opposed to the rate of mutation (a purely physiological parameter), we converted that rate to reflect the difference in generation time, to 0.34 substitutions.site^−1^.Myr^−1^ (95% CI = 0.18–0.54).

We followed the protocol of Cristofari et al.^7^, a development of the protocol of Trucchi et al.^22^, downsampling the data to haploid individuals, and using 50 randomly selected but unlinked RAD loci with 50 haplotypes each, with 3 to 6 polymorphic sites, in addition to 50 randomly selected mitochondrial HVR haplotypes. We specified one independent site model for each locus class (3, 4, 5 or 6 SNPs, and HVR). For each class, we specified a substitution model allowing for invariant sites for the HVR, but not for the short nuclear loci, and for gamma-distributed rate heterogeneity discretised in 4 classes. Transition-transversion ratio *kappa* was linked across nuclear models. All chains were run in duplicate to check for convergence and for a sufficient length to gather ESS > 200 for all parameters, which necessitated 500,000,000 to 1,000,000,000 steps.

Since we parametrised each locus class separately, we expect our model to fit a class-specific substitution rate as a function of the observed number of segregating sites, rather than a common substitution rate. However, as we focus on neutrally evolving regions of the genome, we expect the number of segregating sites to follow a Poisson distribution, of parameter λ equal to the mean number of segregating sites per RAD locus. On a large number of sequences, the expected value E(λ) converges towards the “true” underlying constant mutation rate, multiplied by the total tree length for each locus. Thus, if we fix the tree length, λ becomes an estimator of the substitution rate μ. However, under the EBSP model, the observed number of segregating sites is taken as an estimator of λ, and consequently of the substitution rate μ. Therefore we expect the inferred value of μ for each locus class to be a posterior probability of the “true” substitution rate, conditional on the mean number of segregating sites observed for that class^22^. In order to retrieve the underlying common substitution rate μ, we first fitted a log-linear model to the inferred substitution rates (μ_3_= 0.0159, μ_4_= 0.0218, μ_5_ = 0.0275, μ_6_= 0.0389. Fitted model: intercept *i* = -5.02, slope *s* = 0.292, R^2^=0.997). A Poisson model of parameter μ equal to the mean observed number of segregating sites was a good fit for the empirical distribution of number of segregating sites per locus (λ=1.47, chi-squared test of goodness-of-fit p-value= 0.232). Thus, we extracted μ as *e*^(sλ+i)^ ∼ 1.02e-2 substitutions per site per Myr, or 1.08e-7 substitutions per site per generation.

This rate is ca. twice slower than the one reported by Trucchi et al.^22^ (2.6e-7 subst.site^−1^.generation^−1^), but much faster than the one reported by Li et al.^28^ (8.11e-9 subst.site^−1^.generation^−1^). While the former was not used in Trucchi et al.’s analysis^22^, but rather derived from it, Li et al.’s result ^28^, on the other hand, relies on two exterior and uncertain assumptions: 1) the divergence time between the Emperor and the Adélie penguin is set to ∼23 Myr, which may be a large overestimate Gavryushkina et al.^29^, based on a state-of-the-art total evidence Bayesian analysis, proposes ∼9 Myr instead), and 2) the generation time is taken to be 5 years in both species; however it has been shown to be 16 years in the Emperor penguin^30^, and 6.46 years in the Adélie penguin^27^ - thus 11 years would be a closer (although inaccurate because assuming a single, constant rate) estimate of a common generation time. Applying these corrected estimates to Li et al.’s findings^28^ would give a rate of ∼4.55e-8, which is more than five times faster than proposed, and ca. half our estimate - although this calculation does not take into account the possible rate heterogeneity between lineages, and most importantly the changes in generation time between the *Aptenodytes*/*Pygoscelis* common ancestor and the extant species, which may explain the remaining difference. Generally, the rate of evolution of penguins has been a rather challenging subject, with a wide discrepancy between the paleontological and molecular evidence. While fossil data has been recognised to support a very recent radiation of penguins (about 10 Myr BP^31,32^), molecular data has been interpreted as implying a much more ancient origin (∼45 Myr for Baker et al. ^33^). This molecular-derived radiation has successively been brought to a closer agreement with the fossil evidence by Subramanian et al.^34^ (∼20 Myr) and Gavryushkina et al.^29^ (∼12.5 Myr). The rate that we propose here is in accordance both with the hypothesis of a very fast diversification of the spheniscids, and with the findings of Trucchi et al.^22^.Our reconstruction supports the evidence provided both by the Stairway plot analysis (S2.4) and the PSMC analysis (S2.5), with a fast expansion of the King penguin population in the late Pleistocene, and a stable Emperor penguin population throughout the period (Fig. S4).

The EBSP demographic reconstruction shows only one bottleneck, and places it around 40 Kyr BP - between the two Stairway-inferred bottlenecks. The contrast between the King and the Emperor penguin is maintained, with the Emperor experiencing only a slow and moderate expansion before 100 Kyr BP, and the King going through more diverse demographic events in the late Pleistocene. Our simulation tests (see S2.6) show that, even when two bottlenecks are really present, the EBSP’s expected behaviour is to smooth them out as one single broad population depression (Fig. S7B). Thus, our reconstruction, although with a lower resolution, supports the Stairway-inferred demography. The EBSP’s lower resolution is not surprising, given that it only includes a subset of 50 short loci (*i.e.* 250 to 300 SNPs), where the Stairway plot is using the information from every single genotyped SNP.

**Figure S4.**
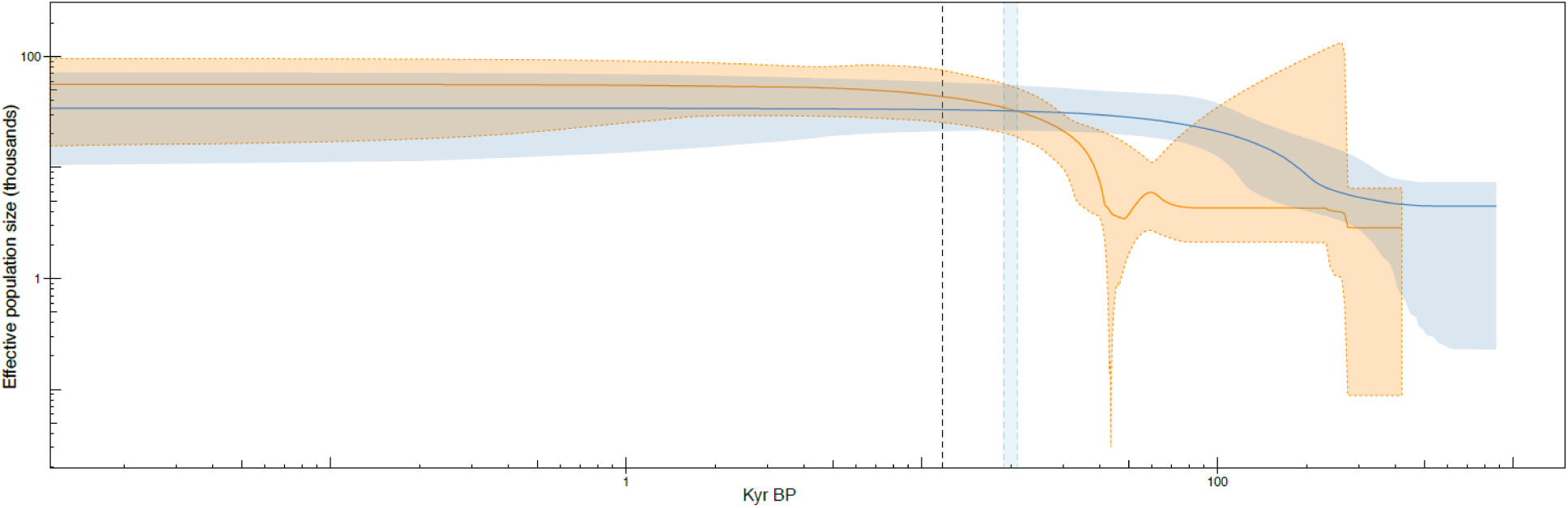
**Extended Bayesian Skyline Plot reconstruction** of past population size changes for the King penguin (orange) and the Emperor penguin (blue). Solid line: median population size; shaded area: 95% confidence interval. Dashed black line: Pleistocene-Holocene boundary. Shaded blue band: last glacial maximum.

### S2.5 Demographic reconstructions III: the Pairwise Sequentially Markovian Coalescent (PSMC’) method on whole-genome sequences

Like the Stairway Plot and the EBSP methods, PSMC ^35,36^ is a model-flexible method, that does not require prior specification of demographic epochs or events. Instead of maximising the composite likelihood of the derived-allele frequency spectrum^24^ or the full likelihood of short, non-recombining sequences^37^, the PSMC algorithm summarises the full ancestral recombination graph through the depth of the most recent coalescence event (time to most recent common ancestor, TMRCA) and total length of singleton branches, as a hidden Markov model in which recombination events mark state changes. It allows for accurate reconstruction of deeper-time demographic events, although it lacks power for more recent time periods in its pairwise form^35,36^. The full Multiple Sequentially Markovian Coalescent approach (MSMC^36^), which has a much improved resolution for recent time periods, relies on the accurate phasing of haplotypes, which unfortunately is not possible in a non-model species, in the absence of a large transmission or population dataset. In order to exploit unphased haplotypes, analysis must be restricted to the pairwise case, as PSMC’. However, since recombination events are treated as a Markovian process along the sequence, it is still possible to increase the likelihood of the reconstruction by concatenating several genomes together, thus increasing the independent sampling of TMRCA. We then selected three high-quality samples for the King penguin, and the Emperor penguin to perform whole-genome re-sequencing. Libraries were prepared with a standard Illumina(c) TruSeq™ PCR-free protocol, and multiplexed on two lanes of a HiSeq 2500 V4 sequencer at the Norwegian Sequencing Center facility, University of Oslo. Reads were mapped to the published Emperor penguin genome^4^ with high success (unique concordant alignment rate, King penguin: ∼86%, Emperor penguin: ∼81%). We retained only longer scaffolds (length ≥ 2 Mb, i.e. 188 scaffolds making up for ∼80% - 1,009,159,582 base pairs - of the total reference length) for the analysis. Concerning each species separately, analysis was run on all three samples simultaneously, with 200 bootstrap replicates. Substitution rate and generation time were defined as above (S2.3-4). Results (Fig. S5) are very similar to the EBSP analysis (S2.4, Fig. S4): the King penguin population grows rapidly in the late Pleistocene, while the Emperor penguin population is mostly stable. However, the resolution of the PSMC’ analysis is low in the recent periods, and the last 4 to 5 time bins exhibit considerable instability when compared across reconstructions ( Fig. S5A), as opposed to older time periods. Thus, the exact timing of the LGM bottleneck is not precisely retrieved for the King penguin: the two-step expansion since the mid-pleistocene (Fig. 1-A in the main text) appears smoothed in one single growth trend. A similar behaviour can be reproduced when simulating data with two bottlenecks in a rapid succession (see S2.6): thus, our PSMC’ analysis is in accordance (although with much lower precision) with our general demography.

**Figure S5.**
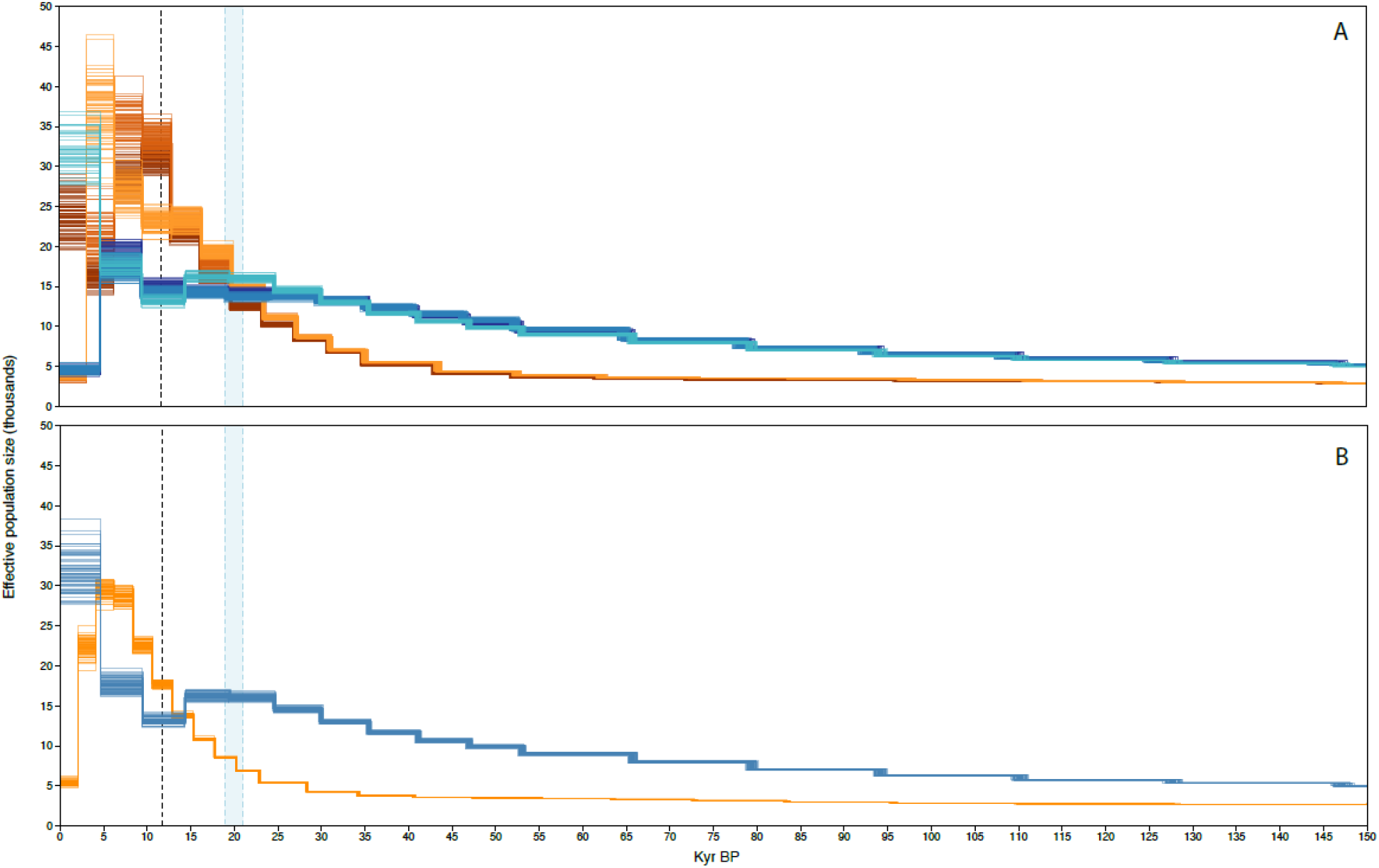
**Pairwise Sequentially Markovian Coalescent reconstruction** of past population size changes for the King penguin (orange) and the Emperor penguin (blue). Each individual line represents one bootstrap replicate. Reconstruction was performed either **A)** for each individual separately as PSMC (each shade represents one individual), or **B)** concatenating genomic data from all three individuals for each species as PSMC’. Dashed black line: Pleistocene-Holocene boundary. Shaded blue band: last glacial maximum.

### S2.6 Comparison of the three demographic inference methods through simulations

In order to assess the consistency of our reconstruction, we simulated genetic data under the Stairway-plot demographic model for the King penguin, and analysed it using all three algorithms (Stairway plot, EBSP, and PSMC’). Data was generated under a sequential Markovian coalescent model, either assuming equal substitution and recombination rates (for the Stairway plot and PSMC’), or 95bp non-recombining haplotypes (for EBSP), using scrm^38^, to match the characteristics of the empirical data, and was either directly converted to an allele-frequency spectrum (for the Stairway plot analysis), or to sequence data, under an HKY model, using seq-gen^39^ (for EBSP). Both the Stairway plot and the PSMC’ approaches rely on bootstrapping, rather than MCMC sampling (as EBSP does), for confidence interval estimation. Whereas the empirical data was bootstrapped directly in a non-parametric way (see S2.4 and S2.5), here, we replicated the full simulation 200 times to estimate the confidence intervals.

**Method I** - The Stairway plot retrieves the principal events in the simulation (Fig. S6A). The main difference lies in the attenuation of the LGM bottleneck, that is mainly visible in the shape of the 95%CI. This is of importance, since it indicates that the Stairway approach may underestimate, rather than overestimate, the bottleneck signal in the data: thus, the bottleneck inferred from the empirical data is likely to be at least as deep as reconstructed. The demographic peak that is visible in the 95%CI at the beginning of the Holocene in our reconstruction from the empirical data ( Fig. 1-A in the main text), on the other hand, although not simulated, is also present in the simulation’s 95%CI. Thus, that secondary peak rather appears to be entirely artefactual.

**Method II** - EBSP on simulated data globally matches the expected demographic history (Fig. S6B), with the true demography nearly entirely included in the EBSP CI95% interval. However, the double bottleneck in our simulated data is smoothed out as one single depression in the reconstruction, that matches neither bottleneck, but rather averages them - although additional complexity is visible in the shape of the lower CI95% interval. When comparing the empirical-data EBSP (Fig. S4), and the simulated reconstructions, CI95% overlap entirely although median effective population size differs, and uncertainty is much larger in the empirical EBSP. Interestingly, however, the empirical run exhibits some features of our simulated model that the simulated-data run fails to retrieve - in particular the low population size during the Llanquihue glacial episode. Due to the low number of SNPs in the loci we include in EBSP analysis, however, less resolution is expected for ancient time periods, so neither the observed discrepancy between simulated and empirical runs, nor the loss of precision compared to the simulated scenario, is surprising.

**Method III** - PSMC’ reconstruction, on the other hand, exhibits a more unexpected behaviour when applied to our data. When assuming equal substitution and recombination rates, none of the bottlenecks is retrieved, but one single bottleneck is inferred instead around 40 Kyr BP, while a large population size peak (absent from our simulation) is inferred in the early holocene (Fig. S6C). Decreasing the recombination rate down to 1/16th of the substitution rate allows us to recover both bottlenecks, yet the artefactual additional population depression remains around 40 Kyr BP, as well as a sharp artefactual population peak after the most ancient bottleneck (Fig. S6D). None of the reconstructions performed on simulated data matches the true demography in a satisfactory way: however, the very recent events on which we focus may be at the limit of the PSMC’ method^35^. It is noteworthy, however, that the empirical PSMC’ inference follows the expected general demographic trend as given by both the Stairway plot analysis and the EBSP analysis, smoothing out both bottlenecks in one single population increase from the early Pleistocene to the late Holocene.

**Fig. S6.**
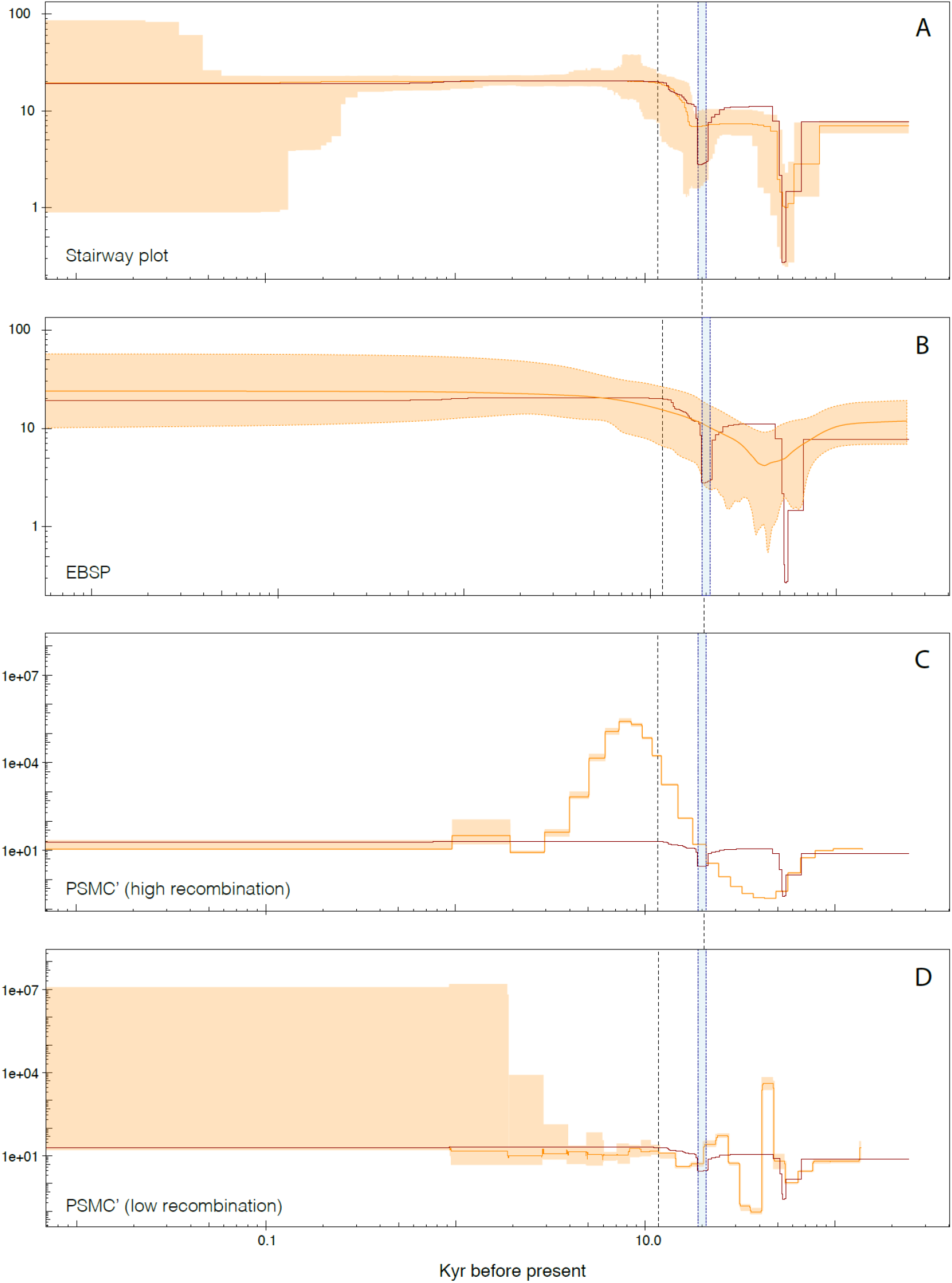
Validation of the demographic reconstructions through simulation. Median effective population size and confidence interval (in thousands of breeders) as a function of time, as reconstructed from simulated data (simulated scenario is represented by the red line on each graph). **A)** Stairway plot reconstruction, **B)** EBSP reconstruction, **C-D)** PSMC’ reconstruction, with either high ( **C**) or low (**D**) recombination rate. Dashed black line: Pleistocene-Holocene boundary. Shaded blue band: last glacial maximum.

## S3 Capture-mark-recapture experiments

In order to verify the hypothesis of high dispersal between colonies, we deployed a capture mark recapture (CMR) experiment on Possession Island, Crozet archipelago, and Ratmanoff beach, Kerguelen archipelago. Ca. 9,832 king penguins were equipped with passive radio-frequency identification (RFID) tags since 1990 on the BDM colony, in Crozet archipelago within the framework of a long-term monitoring program (see Gendner et al.^40^ for details). We deployed mobile detection antennas on all other colonies of Possession Island, as well as on Ratmanoff beach. These antennas have a low detection distance (ca. 60 cm), and are buried in the ground on paths frequented by penguins when they travel in and out of the colony. Each antenna is ∼5 m wide, and records the identification number of any RFID-tagged individual crossing the detection zone. On average, each antenna works for ∼12 hours. Antennas were deployed in the evening, in order to record the activity peak around sunrise. In the current state of development of this system, it is impossible to assess how many individuals (tagged or not) crossed the detection zone during the deployment period: thus, recaptures can only be analysed as presence-absence data, and not as quantitative CMR results. Due to the harsh field conditions, deployments were also in some measure opportunistic; and it is generally impossible to ascertain the status of detected individuals (breeding or moulting), except when their age excluded a breeding attempt. This data, however, provides us precious insights into the behavioural mobility of the species, since antennas were usually located well within the target colonies, and not directly at the seaside: thus, only penguins wandering into a colony distinct from their birth colony were detected.

We performed a total of 28 12-hours deployments during the field seasons 2011-2012, 2012-2013, 2013-2014, and 2014-2015 (Per-colony detections/deployments: CDC: 11/5, PMC: 44/10, JPN: 9/5, MAE:1/1). Out of the 9,832 individuals marked as chicks, an average of 2.3 birds per 12-hour antenna deployment were detected on other colonies of the same island. One anecdotic recapture, in 2014, of a tagged individual born on Crozet in 2009 (and therefore reaching age of first breeding at the time of recapture) also happened on the Ratmanoff beach colony, on Kerguelen archipelago. Although a single event has hardly any statistical value, the Ratmanoff colony counts ∼140,000 breeding pairs^41^, and only two 5m-long antennas were deployed along the beach: thus, this single recapture suggests that dispersal from Crozet to Kerguelen may not be a rare event.

## S4 Palaeoclimate of the Southern Ocean

### S4.1 Definition and constraints of the Antarctic Polar Front

The Southern Ocean is characterised by a strong circular Westerly current that flows uninterrupted by land barriers, the Antarctic Circumpolar Current (ACC). Strong westerly winds generate important northward Ekman transport in the surface water layer, resulting in a convergence of the cold Antarctic surface waters, and warmer Subantarctic surface waters, where the colder southern water masses sink below the northern water mass, at the Antarctic Polar Front (APF). This convergence is compensated by a divergence area, where upwelled deep water masses rise to the surface, creating an intense marine productivity area^42,43,44^. This area is characterised by a steep surface temperature gradient, between 5°C and 3°C^45^.

Generally, a cooling of surface waters in the Southern Ocean is reflected in a northward displacement of the APF, while a warming period brings the APF southward. However, as the APF is defined by the interaction of deep and surface water masses, it is strongly constrained by the sea bottom topography ^45^. Important bathymetric features, such as the Campbell Plateau, the Drake passage, or the Kerguelen plateau, may constrain the position or structure of the APF. In other areas, most importantly the Southern Indian Ocean and Southern Atlantic Ocean, APF displacement is mostly free from bathymetric constraints, and exhibits the largest latitudinal variation^46^.

The Campbell Plateau may be the best studied case of bathymetric constraint on the APF. Both flow models and sediment core evidence showed that the APF remained south of the plateau throughout the Pleistocene, despite important changes in sea surface temperature and frontal positions throughout the Southern Ocean. Whereas the APF is free to move south to greater depths, it is constrained to the North by the sea floor rise^46,47,48^. Similarly, the Drake passage constrains both the northern and southern boundaries of the APF^45,49,50^. Finally, the Kerguelen plateau has been shown to alter the subsurface structure of the front, with its deeper manifestations moving North of the islands, while the surface expressions move South^45,51^. These features, however, are now well modelled in the CMIP5 panel, which has a much improved bathymetric resolution^44^, and the influence of the Drake Passage and Campbell Plateau on the frontal structure is accurately reproduced in our reconstructions (**Fig. 2** in the main text).

### S4.2 Current state of knowledge

There are still considerable uncertainties as to the Pleistocene and Holocene history of the Southern Ocean. Available evidence relies on different types of proxies^46,52,53,54^. **(a)** Ice core data (e.g. EPICA Dome C and Vostok) provide direct evidence for chemical conditions at the core site, and indirect evidence for the oceanic source areas, provided transfer models are accurate enough^55,56^. Parameters derived from ice core evidence mostly covers air temperature, sea ice extent, and marine productivity^56^ **(b)** Benthic sediment core provide more direct evidence for marine conditions (temperature, sea ice cover, productivity) at the core location^52,57^. **(c)** Peat cores and geological evidence on the subantarctic islands and surrounding continental shelf are mostly informative for land ice cover^54^. Taken together, this evidence allows for a general palæoclimatic reconstruction in the Southern Ocean. However, there is still much progress to be done in reconciling the different sources of evidence, as variability amongst core locations (especially benthic sediment cores) is high, and several land-sea coupling mechanisms are still poorly understood^58^. In the current state of knowledge, we can distinguish four major periods in the Southern Ocean late-Pleistocene and Holocene history: (i) Quaternary conditions (59-22 Kyr BP), (ii) Last Glacial Maximum conditions (21-18 Kyr BP), (iii) Pleistocene glacial retreat and early holocene optimum (17-9 Kyr BP), (iv) Holocene hypsithermal and neoglacial conditions (8-0 Kyr BP).

i. ***Quaternary conditions (59-22 Kyr BP)*** were mostly glacial-like, with slow onset of glaciation from ∼35 Kyr BP, and winter sea ice cover reaching as far as ∼56°S in the Pacific. Little is known of land ice throughout the period, as further glaciation obliterated most of the direct evidence.
ii. ***Last Glacial Maximum conditions (21-18 Kyr BP)*** were characterised by extensive land and sea ice cover throughout the Southern Ocean. The Antarctic Polar Front is thought to have moved northward to 40-50°S, a movement associated with a ∼5°C cooling in summer SST^47,52^, although frontal movement is thought to have been constrained by bathymetry south of the Campbell plateau^46,59^. Winter sea ice is also thought to have reached ∼50°S or further northward, or the approximate position of the present-day polar front^60^ (between 47°S in the Atlantic and Indian Oceans, and 57°S in the Pacific Ocean^52^). Marine productivity is thought to have shifted from the Antarctic to the Subantarctic region^54^, while not changing significantly in total biomass^46,56^. Islands of Heard, Crozet, Marion, and the Drake Arc were entirely covered by ice, while Kerguelen and South Georgia may have had ice-free areas. Falklands and Macquarie underwent periglacial conditions^54^. Likely faunal refugia were the currently subtropical islands of Gough, Auckland and Campbell, as well as the Falklands and more generally the Patagonian shelf area^61^.
iii. ***Pleistocene glacial retreat and early holocene optimum (17-9 Kyr BP)*** saw a gradual thawing of most land ice, with contrasting chronologies. Antarctic and subantarctic fronts retreated south to their current location^46^. Sea ice retreated until the Early Holocene climatic optimum (∼11.5-9 Kyr BP), with an episodic increase during the Antarctic Cold Reversal around 14.5 Kyr BP, reaching its current position by ∼10 Kyr BP. Kerguelen and South Georgia archipelagos bear signs of early deglaciation, while Crozet and Marion islands are thought to have carried extensive land ice until the end of the period^54^. The end of the period is marked by a first cold reversal in the Antarctic waters and a short increase in sea ice cover, of unknown extent^62^.
iv. ***Holocene hypsithermal and neoglacial conditions (8-0 Kyr BP)*** were characterised by a warmer climate, similar to historical conditions, interrupted by minor cold reversals. The subantarctic region is ice-free, and the northernmost islands of Gough, Auckland and Campbell are located north of the Subantarctic front^54^. Temperature reaches a maximum around ∼7.5 Kyr BP in the South Pacific^63^. Marine conditions are warm and ice free at ∼50°S until around 6-5 ka BP^62,64^, and temperature drops slightly after ∼3 Kyr BP, although with no change in the glacial landscape. Neoglacial conditions arise after 5 Kyr in East Antarctica, and 3 Kyr in West Antarctica: open water conditions are still prevalent throughout the Southern Ocean, although with possible winter sea ice episodes at 53°S at some periods (∼1-2 Kyr BP).

## S5 Atmosphere-Ocean General Circulation Models (AOGCMs)

### S5.1 AOGCMs choice and multi-model ensemble approach

We used the latest generation of AOGCMs from the IPCC Coupled Model Intercomparison Project Phase 5 (CMIP5)^65^, which represent a significant improvement over CMIP3 in the Southern Ocean^44^. We applied a multi-model ensemble approach, a common improvement over single-model projections, as only the trends present in most models are retained in the final ensemble mean^44^. We selected 15 AOGCMs based on the range of available outputs and their coverage of the Southern Ocean ( Table S1). All model outputs were downloaded from the ESGF nodes (pcmdi9.llnl.gov/). In our study, we used the following variables: Sea Surface Temperature (SST) and Sea-Ice Concentration (SIC). For each variable, we calculated the multi-model ensemble mean and standard deviation using the Climate Data Operators toolset (CDO 2015, available at: http://www.mpimet.mpg.de/cdo).

Reconstructions were performed under Last Glacial Maximum, mid-Holocene, and Historical conditions, and projections according to three Representative Concentration Pathways (rcp) scenarios, the rcp2.6, rcp4.5, and rcp8.5, corresponding respectively to the strong emissions reduction scenario, a moderate emissions profile and the “business-as-usual” scenario. We excluded the rcp6.0 as too few model outputs are available yet.

#### S5.1.1 Sea Surface Temperature (SST)

For palaeoclimate as well as 21^st^ century projections, we followed a protocol similar to that of Péron et al.^66^. The 5°C Sea Surface Temperature (SST) isotherm was used as a diagnostic of the position of the Antarctic polar front (APF) ^45^ where the King penguin is known to forage^66^. The particular breeding cycle of the King penguin makes the constraints on foraging behaviour especially strong during the early chick rearing stage, when the juveniles have not yet reached thermal independence, and need regular feeding while not being able to survive without an adult^67^, which happens around the month of February. This is supported by observed foraging trips, which show a much greater geographic constraint during the month of February^66,68^. Thus, we focused our analysis to the position of the APF in February, as representative of the maximum constraint on foraging trips.

Before using SST outputs derived from AOGCMs, we assessed the accuracy of the representation of the Southern Ocean by comparing each model SST-output for historical runs to satellite-measured SST from december 1981 to december 2005, using the NOAA Optimal Interpolation v2 SST dataset ^69^. Cell-by-cell (1°x1°) linear correlation of SST was assessed and R^2^, slope and intercept were plotted in order to assess the spatial distribution of model departure from observed values.

As modelled SST was generally found warmer than observed SST in the APF zone over the historical period, we followed the correction applied by Péron et al. ^66^. In order to maximise the fit between observed and modelled SST for each archipelago, we defined four oceanic sectors: South Atlantic Ocean (45°W to 18°E), South Indian Ocean (18°E to 80°E), Macquarie (135°E to 180°E), and Falkland region (75°W to 45°W), ranging in latitude from 45°S to 55°S, but extended to 60°S in the Falkland region to account for the higher latitude of the APF around Cape Horn. For each of these sectors, we tested the linear correlation between modelled and observed SST, and we corrected the model value linearly when needed (Table S2). The 5°C SST isotherm was then calculated in GDAL (www.gdal.org), and kilometric distance between each island and the 5°C isotherm was calculated using the OGR Python library. Correctness of our model was assessed through 1) correlating the observed and modelled distances to the 5°C isotherm on the 1981-2005 period and 2) consistency between these distances and published data on King penguin foraging areas.

Foraging range predictions for the historical period closely matched both observed historical SST, and observed foraging distances at most locations: ∼380 km on Crozet (observed: 300-500 km^66^), ∼320 km on Marion (observed: 300 km three decades ago ^70^), ∼20 km in the Kerguelen (observed: 270 km in the APF along the 4 and 5°C isotherms^71,72^ - the APF is reached immediately, but foraging trips extend further in the productivity zone), ∼310 km on Heard (observed: 370 km a decade ago^45^), ∼300 km in South Georgia (observed: 300-600 km over the whole breeding season^72^). Predicted distance for Macquarie Island (∼240 km) is slightly lower than the observed summer range (300-500 km ^73^), however, recorded foraging trajectories meet the APF in the higher-productivity areas on the edge of the Campbell plateau, where upwelling is increased, rather than southward along the shortest route. Finally, the predicted and observed ranges differ most strongly in the Falklands (predicted: ∼640 km and observed: 300-500 km ^74^), a discrepancy explained by the fact the small Falkland population frequently forages on the Patagonian Shelf break, and not directly on the APF^74^. This different behaviour of the Falkland population makes its response to APF displacement more uncertain, as other productivity areas may remain available. However, it seems that the Patagonian Shelf could never sustain a large King penguin population ^75^, and it is sustaining a high, and increasing, anthropogenic pressure from overfishing and climate change^76^. It is therefore unlikely that the Falklands may sustain a significant King penguin population on a centennial time scale.

#### S5.1.2 Winter sea-ice concentration (SIC)

Winter Sea-Ice Concentration (SIC) is known to limit the southward expansion of the King penguin’s breeding range, as the species overwinter breeding cycle makes open-water conditions a requisite throughout the year^67^. Although SIC may still be subject to biases in its representation compared to SST, it has improved since CMIP3 ^77,78,79^. We take the 15% concentration isoline as being representative of the effective sea ice edge^78^. We only consider the sea-ice concentration at their maximum, during the months of august and September. Compared to satellite-derived historical measures from the NOAA Optimal Interpolation dataset, ensemble reconstruction gives a winter sea ice that tends to be more dense than observed values (mean density of sea ice above 15% concentration over the 1981-2005 period: reconstructed 85 ± 20 %; observed 61 ± 22 %, t-test p-value < 2.2e-16), but less extended (reconstructed extent of september SIC > 15% on the 1981- 2005 period occupies 90% of observed SIC > 15% extent), although correlation is strong on a per-cell basis (mean R^2^ = 0.67 ± 0.27). Winter sea ice extent is projected to decrease in all forcing scenarios. While sea ice cover should still be relatively important even at the northern tip of the South Sandwich islands during the last two decades of the century (rcp2.6: 0.26 ±0.058, rcp4.5: 0.22 ±0.044, rcp8.5: 0.045 ±0.040), Bouvet island is projected to become ice-free all year round by 2080 under all forcing scenarios (rcp2.6: 0.058 ±0.037, rcp4.5: 0.028 ±0.024, rcp8.5: 0.00041 ±0.00053). However, sea ice projections may not be quite as reliable as SST projections. Indeed, although geographical distribution is modelled rather accurately, CMIP5 ensemble models fail to reproduce the increase in sea ice extent observed in East Antarctica over the last decades, suggesting that some processes are not yet adequately accounted for in the current models^78^ - in particular, the impact of the influx of fresh meltwater from the Antarctic ice sheet on the extent of winter sea ice may still be widely underestimated. If such a bias exists, however, it underestimates the true extent of sea ice: in that case, the King penguin’s range reduction may be even more drastic than we forecast here, as Bouvet may not be ice-free and suitable for colony establishment by the end of the century.

### S5.2 Uncertainties assessment

Although the use of a multi-model ensemble mean approach is considered to outperform the use of a single climate model, it is also essential to assess the uncertainties related to AOGCMs to evaluate the confidence that can be attached to our results. Outputs of the different AOGCMs may diverge across time and space because they are based on diverse parameterization of natural processes, downscaling approaches, spatial resolutions, etc. To assess the uncertainties associated with our projections, we calculated, for each rcp scenario, the projected foraging distance derived from each climate model separately. We followed the protocol developed by Goberville et al.^80^ by calculating the density distribution of projected foraging distance for each island (i) for the current period (2006-2015), (ii) for the middle of the century (2041-2050) and (iii) for the end of the century (2091-2100) (Extended Data Figure 3). In addition, for the same periods, we also calculated the percentage of models forecasting local King penguin population collapse (February foraging distance > 700 km; Extended Data Figure 3), as proposed by Raybaud et al.^81^. The latitude of the APF, and therefore the duration of the King penguin’s foraging trips, is subject to a high interannual variability, in particular under the influence of the cyclical El Niño Southern Oscillation and Southern Annular Mode, with year-to-year latitudinal fluctuations of up to 200 km^68^. Therefore, we considered that a location had reached its critical foraging distance when foraging distance was higher than 700 km for at least 20% of a consecutive decade.

Variability between models remains relatively high, as has already been observed in previous studies ^80,82^ (Extended Data Figure 3). At all locations, predictions overlap entirely between rcp scenarios for the first decade of our projections, as expected. This is still mostly the case in the middle of the century (2041-2050). Most of the divergence between scenarios appear by the end of the century. This may take the form of (i) a strong divergence of the rcp-8.5 projections as opposed to rcp-4.5 and rcp-2.6 (as in Kerguelen and Bouvet); (ii) an increased dispersion on rcp-8.5 projections (as in Crozet and Prince Edward), or (iii) a more gradual panel of possible outcomes from rcp-2.6 to rcp-8.5 (at most other locations), or (iv) no strong difference between scenarios in Heard Island. This contrast between scenarios is also noticeable when considering the proportion of individual models predicting a local extinction at each time period (Extended Data Figure 4). In the last decade of the century, the ‘business-as-usual” rcp-8.5 scenario stands out compared to the “controlled-emissions” rcp-2.6 and rcp-4.5 in Kerguelen, Crozet, Prince Edward, Bouvet and South Georgia, while all three scenarios make up a gradient in Macquarie, South Sandwich, the Falklands, and Tierra del Fuego. Under rcp-8.5, more than 50% of the models predict extinction in Crozet, Prince Edward and the Falklands by the end of the century, and the difficult position of Kerguelen and Tierra del Fuego is confirmed by the fact that a large proportion of models predict extinction on these islands too. Overall, although inter-model variability remains high, and alternative outcomes are possible, the strong consensus both in the increasing foraging distance trend, and in the actual prediction for local extinction, stress both the very likely nature of the threats upon the Southern Ocean ecosystems under the rcp-8.5 scenario, and the possibility of yet avoiding the most destructive effects of these threats if immediate action allows us to bring greenhouse-gas emissions closer to the rcp-2.6 forcing scenario.

**Table S1.** Ensemble members used in habitat predictions. Model outputs were downloaded from the IPCC archive ( <u>http://www.ipcc-data.org/sim/gcm_monthly/AR5/Reference-Archive.html</u>). Only one ensemble member was used for each model (r1i1p1 whenever available). Not all models outputs for both 21st century and paleoclimate experiments, thus different ensembles were used for LGM, mid-Holocene and 21st century reconstructions.

| Model | Institution | LGM | mid-Holocene | 21st century |
| --- | --- | --- | --- | --- |
| BCC-CSM1 | Beijing Climate Center<br>(China) | N/A | r1i1p1 | r1i1p1 |
| CanESM1 | Canadian Centre for<br>Climate Modelling and<br>Analysis<br>(Canada) | N/A | N/A | r1i1p1 |
| CCSM4 | National Center for<br>Atmospheric Research<br>(USA) | r1i1p1 | r1i1p1 | r1i1p1 |
| CESM1-CAM5 | National Center for<br>Atmospheric Research<br>(USA) | N/A | N/A | r1i1p1 |
| CNRM-CM5 | Centre National de<br>Recherches<br>Météorologiques,<br>Centre Européen de<br>Recherche et de<br>Formation Avancée en<br>Calcul Scientifique<br>(France) | r1i1p1 | r1i1p1 | r1i1p1 |
| CSIRO-Mk3-6-0 | Australian<br>Commonwealth Scientific<br>and Industrial Research<br>Organization (Australia) | N/A | r1i1p1 | N/A |
| EC-EARTH | EC-EARTH consortium<br>published at Irish Centre<br>for High-End Computing<br>(Netherlands/Ireland) | N/A | N/A | r8i1p1 |
| FIO-ESM | The First Institute of<br>Oceanography, SOA<br>(China) | N/A | N/A | r1i1p1 |
| FGOALS-G2 | Institute of Atmospheric<br>Physics, Chinese | r1i1p1 | r1i1p1 | N/A |
|  | Academy of Sciences,<br>and Tsinghua University<br>(China) |  |  |  |
| GFDL-ESM2M | Geophysical Fluid<br>Dynamics Laboratory<br>(USA) | N/A | N/A | r1i1p1 |
| GISS-E2-R | NASA/GISS (Goddard<br>Institute for Space<br>Studies)<br>(USA) | r1i1p1 | r1i1p1 | r1i1p2 |
| HadGEM2-ES | Met Office Hadley Centre<br>(UK) | N/A | r1i1p1 | r1i1p1 |
| IPSL-CM5A-MR | Institut Pierre Simon<br>Laplace<br>(France) | r1i1p1 | r1i1p1 | r1i1p1 |
| MIROC5 | Atmosphere and Ocean<br>Research Institute,<br>National Institute for<br>Environmental Studies,<br>and<br>Japan Agency for Marine-Earth Science and<br>Technology<br>(Japan) | N/A | N/A | r1i1p1 |
| MPI-ESM-MR | Max Planck Institute<br>for Meteorology<br>(Germany) | N/A | N/A | r1i1p1 |
| MPI-ESM-P | Max Planck Institute<br>for Meteorology<br>(Germany) | r1i1p1 | r1i1p1 | N/A |
| MRI-CGCM3 | Meteorological Research<br>Institute<br>(Japan) | r1i1p1 | r1i1p1 | r1i1p1 |
| NorESM1-M | Bjerknes Centre for | N/A | N/A | r1i1p1 |
|  | Climate Research,<br>Norwegian<br>Meteorological Institute<br>(Norway) |  |  |  |

**Table S2.** Correlation of observed and modelled SST in the Southern Ocean. Slope, intercept and correlation coefficient for linear correlation of our ensemble model and observed SST data over the historical period (1981-2005), in four sectors of the Southern Ocean.

| Sector | $R^2$ | Slope | Intercept |
| --- | --- | --- | --- |
| Drake | $0.903 \pm 0.0291$ | $0.683 \pm 0.0472$ | $1.84 \pm 1.19$ |
| South Atlantic | $0.945 \pm 0.0259$ | $0.785 \pm 0.0571$ | $1.84 \pm 1.04$ |
| South Indian | $0.945 \pm 0.0157$ | $0.791 \pm 0.0730$ | $2.27 \pm 0.876$ |
| Macquarie | $0.945 \pm 0.0160$ | $0.805 \pm 0.0687$ | $1.80 \pm 0.705$ |

